# Comparative analysis of genome-encoded viral sequences reveals the evolutionary history of flavivirids (family *Flaviviridae*)

**DOI:** 10.1101/2021.09.19.460981

**Authors:** Connor G. G. Bamford, William M. de Souza, Rhys Parry, Robert J. Gifford

## Abstract

The flavivirids (family *Flaviviridae*) are a group of positive-strand RNA viruses that pose serious risks to human and animal health on a global scale. Here we use flavivirid-derived DNA sequences, identified in animal genomes, to reconstruct the long-term evolutionary history of family *Flaviviridae*. We demonstrate that flavivirids are >100 million years old and show that this timing can be combined with dates inferred from co-phyletic analysis to produce a cohesive overview of their evolution, distribution and diversity wherein the main flavivirid subgroups originate in early animals and broadly co-diverge with major animal phyla. In addition, we reveal evidence that the ‘classical flaviviruses’ of vertebrates, most of which are transmitted via blood-feeding arthropod vectors, originally evolved in hematophagous arachnids and later acquired the capacity to be transmitted by insects. Our findings imply that the biological properties of flavivirids have been acquired gradually over the course of animal evolution. Thus, broad-scale comparative analysis will likely reveal fundamental insights into their biology. We therefore published our results via an open, extensible, database (Flavivirid-GLUE), which we constructed to facilitate the wider utilisation of genomic data and evolution-related domain knowledge in flavivirid research.

## INTRODUCTION

The flavivirids (family *Flaviviridae*) are an important group of RNA viruses incorporating numerous pathogens of humans and animals. Currently, four genera are recognised within the family: *Pegivirus, Pestivirus*, *Hepacivirus* and *Flavivirus* [1]. While pegiviruses are not known to be associated with disease, pestiviruses cause serious illness in domestic ungulates such as cattle and pigs [2], and the *Hepacivirus* genus includes the bloodborne hepatitis C virus (HCV), a major cause of chronic liver disease in human populations throughout the world [3, 4]. Moreover, the genus *Flavivirus* includes viruses that are transmitted between vertebrates via blood-feeding arthropod vectors (e.g., mosquitoes, ticks) and cause large-scale outbreaks resulting in millions of human infections every year - e.g., yellow fever virus (YFV), dengue viruses 1-4 (DENV1-4), Zika virus.

The *Pegi-, Pesti-*, *Hepaci-* and *Flavivirus* genera contain viruses with monopartite genomes ∼10 kilobases (Kb) in length and encoding one or more large polyproteins that are co- and post-translationally cleaved to generate mature virus proteins. The structural proteins of the virion - capsid (C), premembrane (prM) and envelope (E) - are encoded toward the 5’ end of the genome, while genes encoding non-structural (NS) proteins are located further downstream [5]. However, a diverse variety of novel ‘flavivirus-like’ viruses (flavivirids) have been identified in recent years and these viruses – most of which were identified in invertebrates and have yet to be incorporated into official taxonomy - exhibit a much greater range of variation in genome structure, with genome lengths ranging up to 20Kb [6–10]. Furthermore, one novel group – ‘jingmenvirus’ – comprises viruses with genomes that are multipartite rather than monopartite [11]. Some tick-associated jingmenviruses have been linked with disease in humans [12, 13].

To prevent the spread of pathogenic viruses, it is helpful to understand their evolutionary history in as much detail as possible, because this can often provide crucial insights into virus biology and host-virus relationships [14]. Flavivirids are a taxonomically diverse group that have been extensively examined using comparative approaches, revealing uniquely clear correlations between phylogenetic relationships and ecological characteristics [15–18]. The current, rapid accumulation of genome sequence data from novel flavivirids offers unprecedented opportunities to build on these comparative studies. Presently, however, there are two major obstacles to efficient use of flavivirid genome data.

Firstly, there is a general lack of re-use and reproducibility among comparative analysis of virus genomes, especially when deeper evolutionary relationships are being examined [19]. These analyses typically entail the assembly of complex data sets comprised of virus sequences, multiple sequence alignments (MSAs) and phylogenies linked to other, diverse kinds of data (e.g., spatiotemporal coordinates, taxonomy, immunity-related information). In theory, these data sets could be re-used across a wide range of analysis contents while also being collaboratively developed and refined by multiple contributors. This would likely accelerate knowledge discovery and expedite the development of expert systems utilising virus genome data. Unfortunately, however, such practices remain challenging to implement in practice, largely due to a lack of appropriate tools and data standards [20].

Secondly – knowledge of the appropriate evolutionary timescale is critically lacking. So far, most studies of flavivirid evolution have proposed relatively short timelines in which individual genera and sub-groups emerge within the last 10-100 thousand years [15–17, 21]. However, these studies were based on viruses sampled from a relatively restricted range of hosts. By contrast, recent studies utilising metagenomic techniques to sample flavivirid diversity across a broader range of animal species have prompted suggestions of a much longer timeline extending over hundreds of millions of years [7]. Furthermore, for many RNA virus families, robust evidence for ancient origins has come in the form of *endogenous viral elements* (EVEs) - virus-derived sequences found within eukaryotic genomes [22]. These sequences are thought to originate when infection of germline cells leads to virus-derived cDNA being incorporated into chromosomal DNA so that integrated viral genes are not only inherited as host alleles, but also persist in the gene pool over many generations until they are genetically fixed (i.e., reach a frequency of 100% in the species gene pool). Genome comparisons show that EVE loci are often present as orthologs in closely related host species, establishing their ancient origins (since they were incorporated into the host germline prior to species divergence). EVEs derived from flavivirids have been identified in a handful of arthropod species [8, 23–25], but are relatively uncommon compared to EVEs derived from other RNA virus families [26]. Partly reflecting this scarcity, robust calibrations of the long-term evolutionary timeline of flavivirids are lacking.

In this study, we calibrate the long-term evolutionary timeline of flavivirids, making extensive use of EVEs. In addition, we construct a cross-platform, interactive database called ‘Flavivirid-GLUE’, which we use to capture the evolution-related domain knowledge generated in our study in a way that facilitates downstream use.

## RESULTS

### Creation of open resources for comparative genomic analysis of flavivirids

We previously developed a software framework called GLUE (“<u>G</u>enes <u>L</u>inked by <u>U</u>nderlying <u>E</u>volution”) [27]. Here, we used GLUE to create Flavivirid-GLUE [28] - a flexible, extensible, and openly accessible resource for comparative analysis of flavivirid genomes (**Fig. S1a-b**). The Flavivirid-GLUE project includes: (i) a set of 237 reference genome sequences each representing a distinct flavivirid species and linked to isolate-associated data (**Table S1**) (ii) a standardized set of 81 flavivirid genome features (**Table S2**); (iii) genome annotations specifying the coordinates of genome features within selected ‘master’ reference genome sequences (**Table S3**); (iv) a set of hierarchically arranged MSAs constructed to represent distinct taxonomic ranks within the family *Flaviviridae* (**Table 1**, **Fig. 1**).

**Figure 1.**
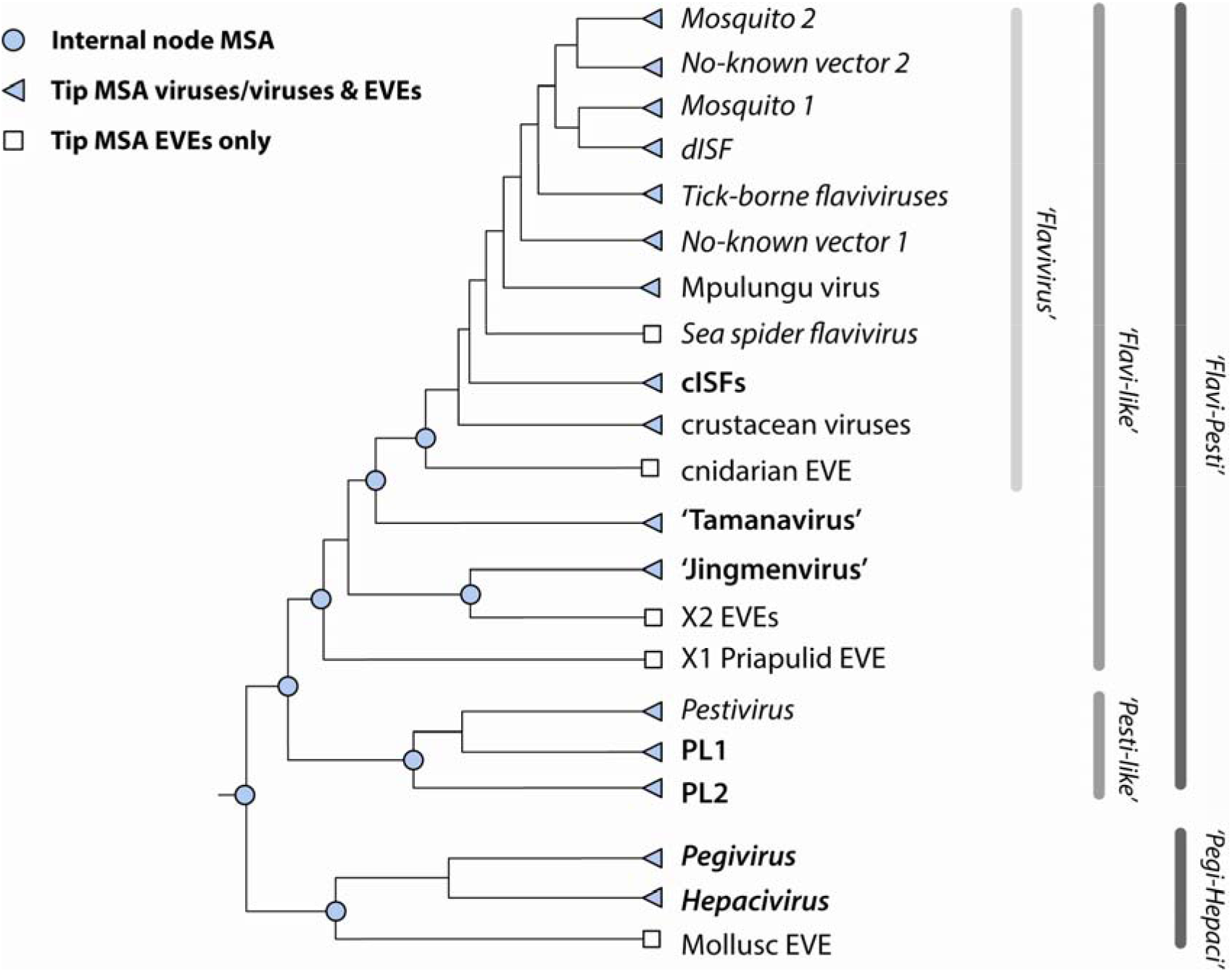
The multiple sequence alignment hierarchy in Flavivirid-GLUE. The GLUE software framework defines a ‘constrained alignment tree’ data structure comprising a set of alignments that are hierarchically linked to reflect taxonomic relationships [27]. To enable sequence comparisons across the entire *Flaviviridae*, we implemented a constrained alignment tree data structure in *Flavivirid*-GLUE, as shown in the cladogram above. Numbers shown adjacent to nodes correspond to rows in **Table 1. Abbreviations**: cISF = classical insect-specific flaviviruses; NKV = no known vector; Jingmen = Jingmen tick virus =; PL = ‘pestivirus-like’; MBFV = mosquito borne flavivirus; MSA = multiple sequence alignment.

**Table 1.** Comprehensive mapping of flavivirid homology via hierarchically linked multiple sequence alignments. ***^a^*** Numbers correspond to labelled nodes in **Fig. 1**. ***^b^***The parent multiple sequence alignment (MSA) in the hierarchy. ***^c^*** Children of the MSA in the hierarchy. ***^d^*** Reference sequence that constrains the genomic co-ordinate coordinate space in the MSA. ***^e^*** Percentage of the constraining reference genome spanned by the MSA. Phylogenies constructed for each of these alignment partitions are available in Flavivirid-GLUE [28]. ***^f^***Counts reflect the number of consensus EVE sequences included in the alignment – consensuses are linked to child MSAs that contain the sequences of all individual EVE loci used to create the consensus.

| # <sup>a</sup> | Taxonomic scope | Name | Parent <sup>b</sup> | Children <sup>c</sup> | Constraining reference <sup>d</sup> | Genome Coverage <sup>e</sup> | Virus count | EVE count <sup>f</sup> |
| --- | --- | --- | --- | --- | --- | --- | --- | --- |
| **Family** |  |  |  |  |  |  |  |  |
| 1 | Flaviviridae-like | Flaviviridae | none | 2 | YFV | ~6% | 2 |  |
| **Major lineage** |  |  |  |  |  |  |  |  |
| 2 | Hepaci/Pegi-like viruses | HepaPegi | Flaviviridae | 2 | BVDV1 | ~60% | 2 | 1 |
| 3 | Flavi/Pesti-like | FlaviPesti | Flaviviridae | 2 | YFV | ~80% | 2 | 1 |
| 4 | Flavi-like | FlaviLike | FlaviPesti | 2 | YFV | ~35% | 2 |  |
| 5 | Pesti-like | PestiLike | FlaviPesti | 3 | BVDV1 | ~60% | 3 |  |
| **Minor lineage** |  |  |  |  |  |  |  |  |
| 6 | Flavi-Tamana-like | FlaviTamana | FlaviLike | 2 | YFV | ~93% |  |  |
| **Genus** |  |  |  |  |  |  |  |  |
| 7 | *Flavivirus* genus | Flavivirus | FlaviTamana | 11 | YFV |  | 8 | 3 |
| 8 | *Hepacivirus* genus* | Hepacivirus | HepaPegi | 0 | HCV | ~90% | 14 |  |
| 9 | *Pegivirus* genus* | Pegivirus | HepaPegi | 0 | HPgV2 | ~60% | 9 |  |
| 10 | Insect pesti-like* | PL2 | PestiLike | 0 | SLV2 | ~73% | 6 | 7 |
| 11 | SCNV+arthropod* | PL1 | PestiLike | 0 | SCNV5 | ~72% | 8 |  |
| 12 | *Pestivirus* genus* | Pestivirus | PestiLike | 0 | BVDV1 | ~88% | 11 |  |
| 13 | X2-derived EVEs | X2 | FlaviLike | 0 |  | NS5 |  | 9 |
| 14 | 'Jingmenvirus' seg 1 | Jingmen_1 | FlaviLike | 0 | JMTV Seg1 | Segment 1 | 7 | 2 |
| 14 | 'Jingmenvirus' seg 3 | Jingmen_3 | FlaviLike | 0 | JMTV Seg3 | Segment 3 | 4 | 1 |
| 15 | 'Tamanavirus' | 'Tamanavirus' | FlaviTamana | 0 | TABV | 100% | 6 | 5 |
| **Subgenus** |  |  |  |  |  |  |  |  |
| 16 | Crustacean | crustacean | Flavivirus | 0 |  | 100%Δ-genome | 2 |  |
| 17 | cISF | cISF | Flavivirus | 0 | KRV | 100% | 14 | 6 |
| 18 | NKV1 | NKV2 | Flavivirus | 0 | APOIV | 100% | 6 |  |
| 19 | Tick | tick | Flavivirus | 0 | POWV | ~91% | 15 |  |
| 20 | dISF | dISF | Flavivirus | 0 | LAMV | ~95% | 9 |  |
| 21 | Mosquito1 | mosquito-1 | Flavivirus | 0 | DEN1 | ~97% | 35 |  |
| 22 | NKV2 | NKV2 | Flavivirus | 0 | SOKV | 100% | 3 |  |
| 23 | Mosquito2 | mosquito-2 | Flavivirus | 0 | YFV | ~93% | 14 |  |
| **Totals** |  |  |  |  |  |  | **159** | **35** |

We used hierarchically linked MSAs to enable standardised genome sequence comparisons across the entire *Flaviviridae* family (i.e., both within and between taxonomic ranks). For each taxonomic rank represented in the project, one reference genome was selected as the constraining ‘master’ reference that defines the genomic coordinate space. Within the MSA hierarchy, each MSA is linked to its child and/or parent MSAs via our chosen set of references. MSAs representing internal nodes (see **Table 1**) contain only master reference sequences but can be recursively populated with all taxa contained in child alignments via GLUE’s command layer [27]. Importantly, the use of an MSA hierarchy simplifies analysis of novel taxa (since new sequences only need be aligned to the most closely related reference genome to be aligned with all other taxa included in the MSA hierarchy).

Instantiation of the Flavivirid-GLUE project (**Fig. S2**) generates a relational database that contains the data items required for comparative analysis of flavivirids and represents the semantic links between them. This allows comparative analyses to be implemented in a standardised, reproducible way, wherein GLUE’s command layer is used to coordinate interactions between the Flavivirid-GLUE database and bioinformatics software tools (e.g., see **Fig. S3-S4**). Flavivirid-GLUE can be installed on all commonly-used computing platforms, and is fully containerised via Docker [29]. Hosting in an openly accessible online version control system (GitHub) provides a platform for coordinating ongoing development of the resource – e.g., incorporation of additional taxa and genome annotations - following practices established in the software industry (**Fig. S1c**) [30].

### Mapping the distribution of flavivirid-derived DNA in animal genomes

To identify flavivirid-derived EVEs, we performed systematic *in silico* screening of whole genome sequence data representing 1075 animal species. This led to the identification of 374 EVE loci in 36 animal species (**Table 2**, [28]). We reconstructed consensus sequences representing fragments of the genomes of (presumably) extinct flavivirids, utilising EVE sequences putatively derived from a single germline incorporation event (i.e., orthologs, fragments, duplicates) (**Fig. 2**, **Fig. S5**). EVE consensus sequences, MSAs, and all EVE-associated metadata were incorporated into Flavivirid-GLUE [28].

**Figure 2.**
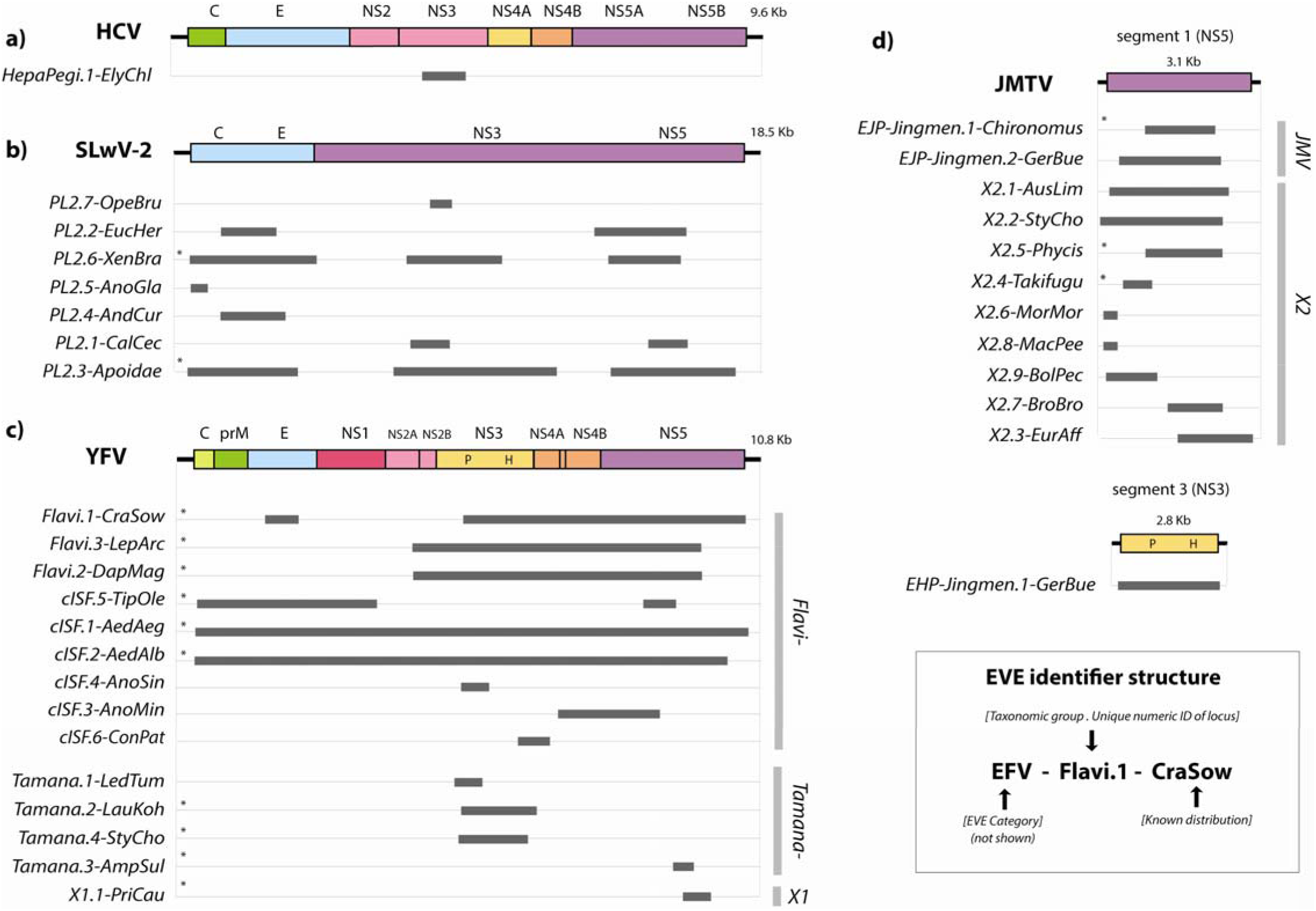
Genome structures of endogenous flaviviral elements. Schematic diagrams showing the genomic regions represented by flavivirid-derived endogenous viral element (EVE) sequences. **(a)** Pegi-hepacivirus-like elements are shown relative to hepatitis C virus (HCV); **(b)** elements derived from the ‘Pestivirus-like 2’ (PL2) lineage of viruses shown relative to Shuangao lacewing virus 2 (SLwV-2); **(c)** Elements derived from the *Flavivirus* genus, the ‘tamanaviruses’, and the X1 lineage shown relative to Dengue virus type 1 (DENV-1); **(d)** ‘jingmenvirus’-derived EFVs shown relative to Jingmen tick virus (JMTV). Homologous regions represented by EFV sequences are shown as grey horizontal bars. Bars to the right show taxonomic groups. EVE identifiers (IDs) are shown to the left. EVE IDs were constructed as indicated in the key, following a convention established for endogenous retroviruses [70]. IDs have three components - the first is the classifier ‘EFV’ ‘endogenous flaviviral element’ (can be dropped when implied by context). The second comprises two subcomponents separated by a period; (i) the name of the taxonomic group of viruses from which the EFV is thought to derive; (ii) a numeric ID that uniquely identifies the integration locus. The third component describes the known taxonomic distribution of orthologous copies of the element among host species. For EVEs derived from the ‘jingmenvirus’ lineage, which contains viruses with multipartite genomes, we used classifiers that specify the gene from which the EVE is derived, in line with conventions established for EVEs derived from segmented viruses [71]– endogenous ‘jingmenvirus’ helicase (EJH) and endogenous ‘jingmenvirus’ polymerase (EJP). **Abbreviations**: Kb = kilobase. X1= unclassified flavivirus-like lineage X1. X2 = unclassified flavivirus-like lineage X2. C = Capsid, prM = Pre-Membrane, E = Envelope, NS = Non-Structural protein, P = Protease, H = Helicase. HP = ‘pegi-hepaci’. **\*** Indicates consensus sequence.

**Table 2.**
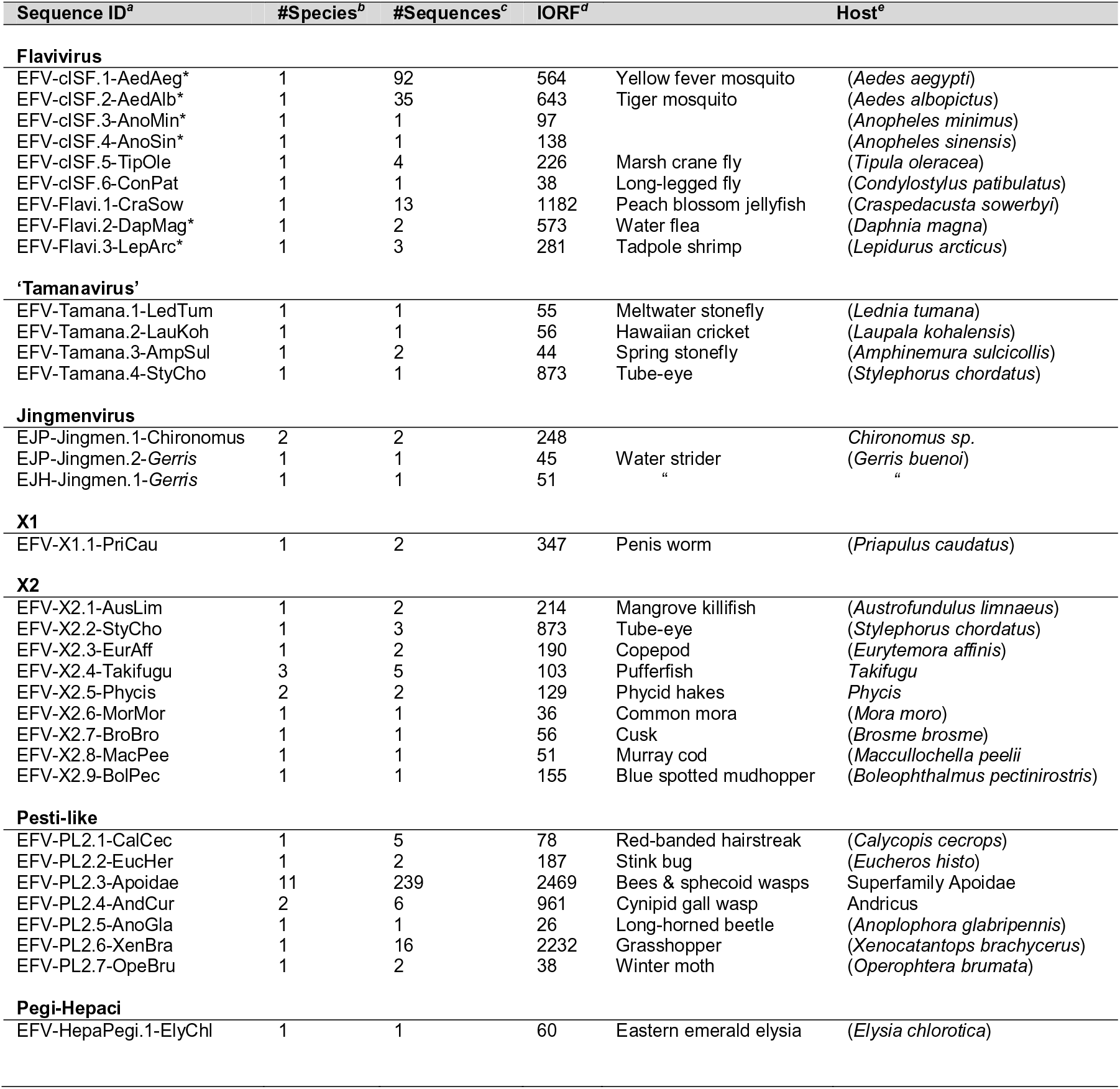
Flavivirid-derived endogenous viral elements. ***^a^*** Flavivirid-derived EVEs have been assigned standard IDs based on conventions established for endogenous retroviruses, wherein information about virus taxonomy and locus orthology are incorporated into the ID itself [70]. The ID comprises of three elements separated by hyphens. For most EVEs characterised here, the first (i.e., leftmost) element is the classifier ‘endogenous flavivirid’ (EFV). However, for ‘jingmenviruses’ the classifier component of the ID also specifies the gene it derives from (EJP=endogenous jingmenvirus polymerase; EJH=endogenous jingmenvirus helicase), following conventions established for multipartite viruses and EVEs derived from mRNA sources [71]. The second ID element comprises two subcomponents separated by a period – the first defines the taxonomic position of the EVE in relation to established *Flaviviridae* taxonomy, the second is a numeric ID that uniquely represents an EVE locus. The third ID component defines the known distribution of orthologous insertions among host species. If it is only known from a single species a shortened version of the Latin binomial species name is used. ***^b^*** Number of species in which EVE locus was identified. ***^c^*** Number of sequences (i.e., distinct insertions) derived from this EVE that were identified via *in silico* screening. ***^e^*** Host species or species groups. Asterisks indicate EFVs loci or lineages that have been reported previously. Abbreviations: L-ORF=Longest open reading frame; PL2=Pesti-like 2; cISF=classical insect-specific flaviviruses.

### All major flavivirid lineages are represented in the host germline

We reconstructed the evolutionary relationships between EVEs and contemporary flavivirids using maximum likelihood approaches (**Fig. 3**, **Fig. S6**). Bootstrapped phylogenetic trees were reconstructed from MSAs representing a range of taxonomic ranks within the *Flaviviridae* (**Table 1**), utilising EVE sequences often contain mutations acquired through neutral reconstructed phylogenies both with MSAs containing virus sequences only, and MSAs incorporating both virus and EVE sequences [28]. Consistent with official taxonomy and previously published studies [1, 18] phylogenetic reconstructions split flavivirids into two major lineages - ‘pegi-hepaci’ and ‘flavi-pesti’ – each of which contains several well-supported subgroups [10] (**Fig. 3a-e, Fig. S3**). However, a divergent, flavivirid-derived EVE identified in the genome of a priapulid worm (*Priapulus caudatus*: Lamarck, 1816) may represent a third sub-lineage (**Fig. 3b**).

**Figure 3.**
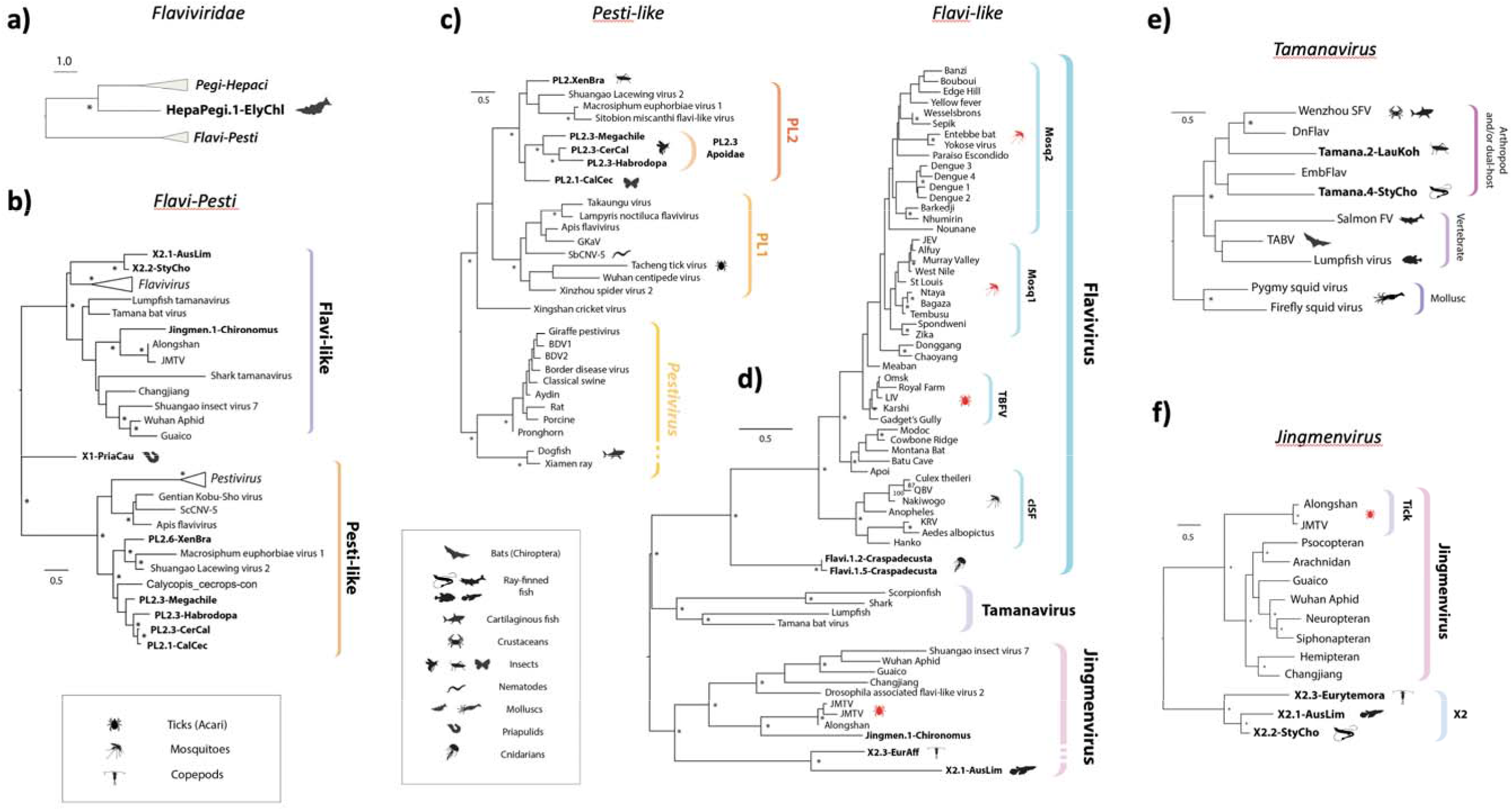
Evolutionary relationships between modern and ancient flavivirids. Bootstrapped maximum likelihood phylogenies (1000 replicates), reconstructed for viruses and EFVs across a range of taxonomic ranks, as follows: **(a)** Two major lineages within family *Flaviviridae* (‘flavi-pesti’ and ‘pegi-hepaci’) showing placement of Hepacivirus-derived EVE (122 amino acid (aa) residues in multiple sequence alignment (MSA) spanning conserved regions in NS3, substitution model=RTREV); **(b)** ‘Flavi-pesti’ lineage (MSA spanning 99 aa residues in NS5, substitution model=BLOSUM62); **(c)** ‘Pesti-like’ lineage (104 aa residues in NS5, substitution model=RTREV); **(d)** ‘Flavi-like’ lineage (MSA spanning 179 aa residues in NS5, substitution model=LG likelihood); **(e)** ‘Tamanavirus’ (MSA spanning 292 aa residues in NS5, substitution model=BLOSUM62); **(f)** ‘Jingmenvirus’ and related lineages (MSA spanning 727 residues in NS5, substitution model=LG likelihood). EFV names are shown in bold. Only EFVs that had sufficient coverage (>50% of total MSA length) were included in the analysis. Triangular terminal branches indicate collapsed clades containing multiple taxa. Asterisks indicate bootstrap support ≥70% (1000 replicates). The scale bar indicates evolutionary distance in substitutions per site. Brackets to the right indicate genera and sub-lineages. All trees are midpoint rooted for display purposes. Host and known/suspected vector associations are indicated by animal silhouettes as shown in the key. **Abbreviations**: cISF = classical insect-specific flaviviruses. NKV = no known vector. Jingmen = Jingmen tick virus =. PL = ‘pesti-like’; MBFV = Mosquito borne flavivirus.

Phylogenetic reconstructions revealed that EVEs derived from a broad range of flavivirus lineages and subgroups are represented in the genomic ‘fossil record’. Only one EVE derived from the ‘pegi-hepaci’ lineage was identified. However, it occurs in a marine mollusc - the Eastern emerald elysia (*Elysia chlorotica*: Gould, 1870) - demonstrating that the host range of this major flavivirid group extends to invertebrates (**Fig. 3a**).

The majority of flavivirid EVEs derived from the ‘flavi-pesti’ lineage, which is comprised of robustly supported ‘flavi-like’ and ‘pesti-like’ clades. Our approach to phylogenetic reconstruction, which entails reconstructing separate phylogenies for distinct taxonomic ranks (see **Table 1**), supports a clean division of ‘flavi-like’ viruses into three monophyletic clades (**Fig. 3d**) corresponding to genus *Flavivirus*, the ‘jingmenviruses’, and a clade of viruses related to Tamana bat virus (TABV) which we here refer to as ‘tamanaviruses’. We identified several EVEs that grouped robustly within the diversity of contemporary ‘tamanavirus’ and ‘jingmenvirus’ isolates. We also identified EVEs derived from a more distantly related, ‘jingmenvirus-like’ lineage – here labelled X2 – with no known contemporary representatives (**Fig. 3f**). Notably, we identified X2-derived EVEs in a temorid copepod (*Eurytemora affinis*: Poppe, 1880) as well as in multiple actinopterygiid fish, indicating that their host range encompasses both vertebrate and arthropod species (**Fig. 3f, Table 1**). The copepod element exhibited internal duplications and rearrangements typical of EVEs identified in piRNA clusters (data not shown) [31].

The ‘pesti-like’ lineage contains two robustly-supported clades – one comprised of vertebrate viruses (including the canonical members of genus *Pestivirus*) while the other contains a diverse assortment of invertebrate-associated, ‘large-genome flavivirids’ [10]. The invertebrate clade contains two well-supported subclades, here labelled ‘PL1’ and ‘PL2’. The ‘PL2’ clade contains EVEs in addition to viruses (**Fig. 3c**).

### Germline incorporation of flavivirid DNA is relatively rare

While flavivirid-derived EVEs were only identified in a small proportion of the animal species we screened, they occur at relatively high copy number in the germline of some insect groups, including mosquitoes of genus *Aedes* (Meigen, 1818) as well as bees and sphecoid wasps (superfamily Apoidea: Latreille, 1802) (**Table 1**).

In the *Aedes* germline, flavivirid EVEs are closely related to contemporary flaviviruses – specifically the ‘classical insect-specific’ flaviviruses’ (cISFs). Numerous, distinct loci occur, largely representing distinct regions of the flavivirus genome (**Fig. S5**). However, where they do span homologous regions of the flavivirus genome loci are highly related (i.e., <1% nucleotide sequence divergence) and are frequently arranged as tandemly repeating arrays (data not shown), suggesting recent, intra-germline amplification.

Among *Apoidea* EVEs, copy number was dramatically inflated in certain species, such as the spurred ceratina (*Ceratina calcarata*: Robertson, 1900), which contains at least 86 distinct flavivirid-derived EVE sequences in its published genome sequence. However, confident estimates of EVE copy number in *Apoidea* species could not be obtained, due to the limitations of current genome assemblies.

### Major flavivirid lineages originated in the distant evolutionary past

We used a range approaches to calibrate the evolutionary timeline of flavirids (**Table 3**). Calibrations based on identification of orthologous EVEs were obtained for ‘jingmenvirus’-derived EVEs found in midges (*Chironomus*: Meigen, 1803), X2-derived EVEs in ray-finned fish (Class Actinopterygii: Klein, 1885), and a PL2-derived EVE in superfamily Apoidea. Orthologous, PL2-derived EVEs were identified in Apoidea species estimated to have diverged >100 million years ago (Mya) (**Fig. S7**). We also derived age estimates ranging between 3-62 Mya for pairs of putatively duplicated EVE sequences based on the assumption a neutral molecular clock (**Table 3**). These calibrations, combined with the identification of flavivirid-derived EVEs in basal animal lineages such as cnidarians and priapulids, suggested that flavivirids could in fact have truly primordial origins in multicellular animals. Such an extended evolutionary timeline would be consistent with other recent data supporting the ancient origins of RNA virus families [7, 32].

**Table 3.**
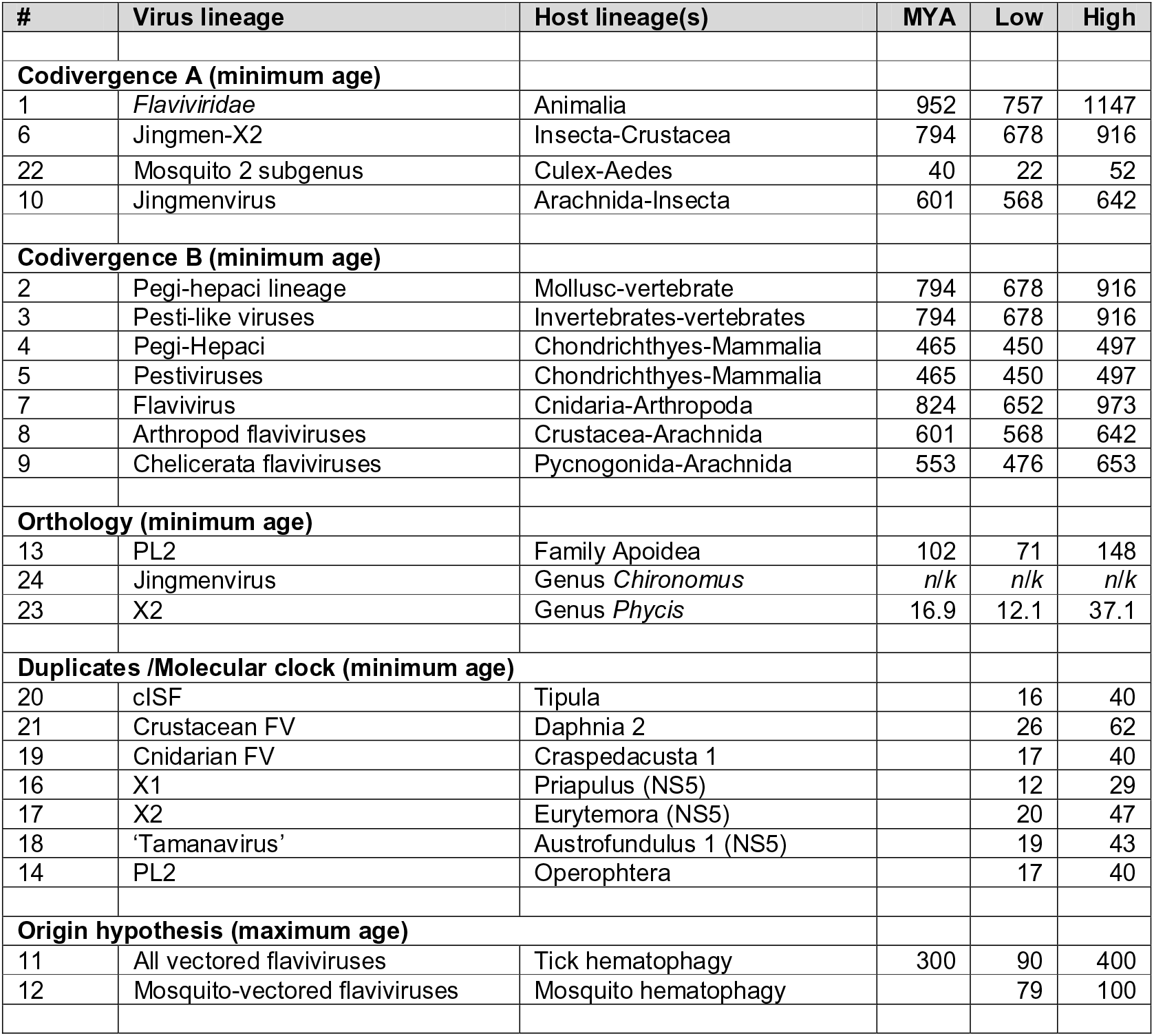
Dates and age estimates used to calibrate flavivirid evolution. Node numbers correspond to those shown in **Fig. 5c**. Hematophagy estimates obtained from [43]. *Culex*-*Aedes* divergence date obtained from [73]. All other divergence data estimates were obtained from TimeTree [72].

Horizontal transfer of flavivirids between distantly related hosts has clearly occurred – most likely in association with parasitism [33, 34] - but when phylogenies are considered in the light of an evolutionary timescale extending back to the origin of multicellular animals (i.e., ∼500-800 Mya), a credible argument can be made for codivergence being the more common mode of evolution, at least where higher taxonomic ranks are concerned. To investigate this, we compared host and virus phylogenies. Scope for comparisons was limited due to sparse data and relatively narrow sampling across host species groups (with most flavivirids isolated from arthropods and vertebrates). We identified several host and virus clades in which the branching relationships and divergence times among animal lineages are correlated with the topology and branch lengths found in virus phylogenies (**Fig. 4, Fig. S8**). However, we were required to make strong assumptions in each case, particularly regarding the rooting of virus trees (see **Fig. 4** legend), and these deeper calibrations should therefore be taken as tentative.

**Figure 4.**
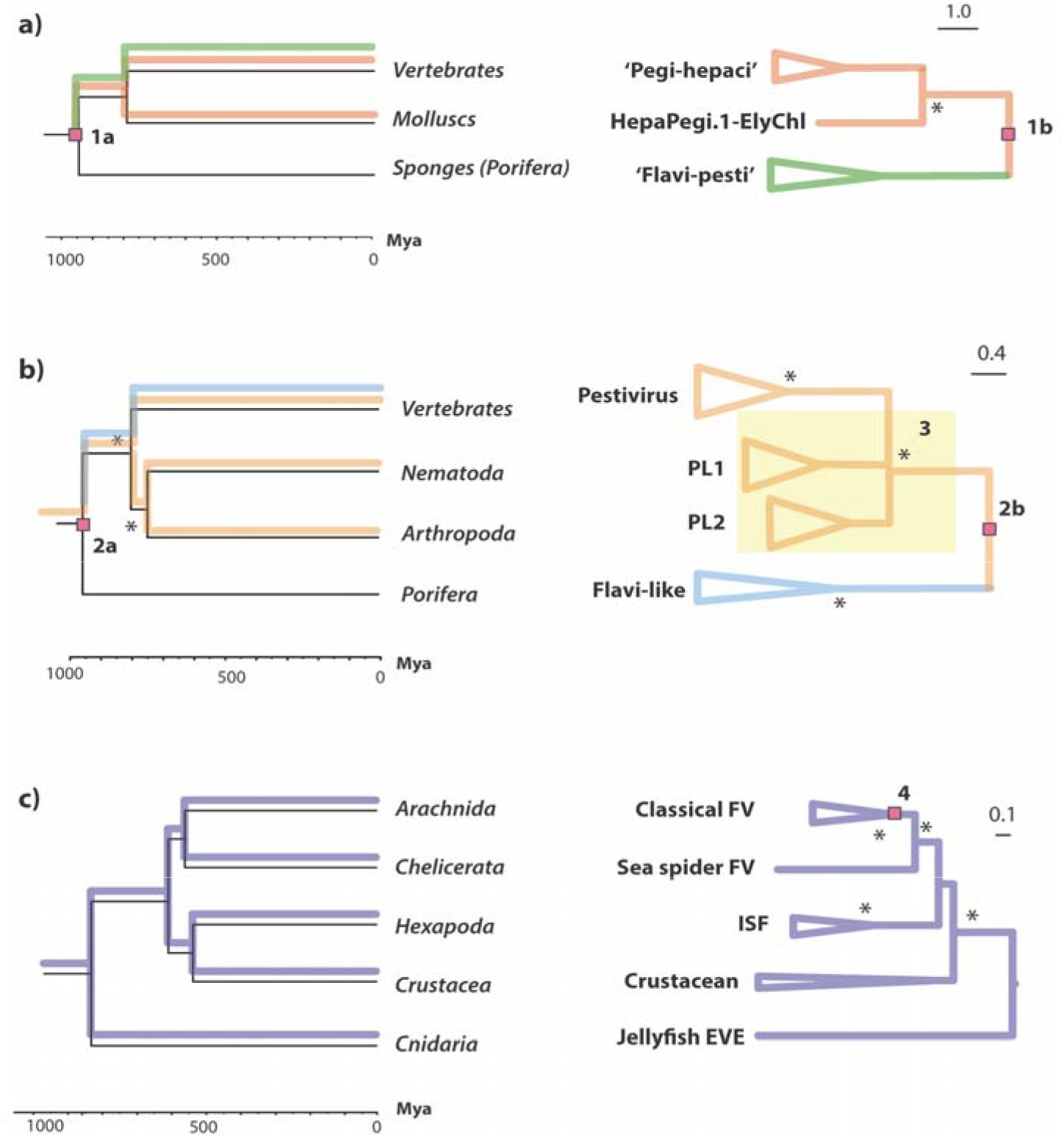
Putative codivergence of flavivirid groups and host phyla. ‘Tanglegrams’ illustrating matching topologies in animal (left) and flavivirid (right) phylogenies. Putative tracking of host lineages by viral lineages is indicated on the host phylogeny. Clades within which the branch lengths of virus phylogenies are correlated with divergence times in host animal lineages: the invertebrate and vertebrate splits in the (**a**) ‘pegi-hepaci’ lineage (R^2^=0.5); (**b**) ‘pesti-like’ lineage (R^2^=0.7); (**c**) the cnidarian-arthropod, chelicerate-hexapoda, and crustacean-insect splits in the *Flavivirus* genus (R^2^=0.3). Note that, due to limited data, all comparisons rely on strong assumptions regarding virus phylogenies, indicated in the figure by numbers, as follows: (1) and (2) midpoint rooting was used, and the deepest divergence in the virus tree was approximately calibrated using the divergence date of porifera (the most basal animal lineage) in line our hypothesis that major flavivirid lineages originated in early metazoans; (3) the splits between nematode and arthropod viruses in “pesti-like 1” (PL1) and “pesti-like 2” (PL2) clades – indicated by the yellow square - were poorly resolved in our phylogenies; (4) midpoint rooting was used, and it is assumed that the arthropod-borne classical flaviviruses originated in arachnids, as proposed in Fig 5a. Abbreviations: Mya=Million years ago.

### Arthropod-vectored flaviviruses likely emerged from an arachnid source

The *Flavivirus* genus includes viruses that are transmitted among vertebrate hosts by arthropod vectors, as well as viruses that exclusively infect arthropods [35] and others that have been identified in vertebrates but have no known arthropod vector [36]. The long history of association between flavivirids and their hosts implied by our investigation suggests the largely vector-borne ‘classical flaviviruses’ could have emerged in association with the evolution of hematophagy (blood-feeding) in arthropods. Phylogenetic reconstructions using either NS5 (**Fig. 3a**) or NS3 (**Fig. S9**) show that flaviviruses exclusively associated with insects (Class Insecta: Linnaeus, 1758) are robustly separated from those that infect vertebrates by viruses identified in crustaceans (Subphylum Crustacea: Brünnich, 1772), sea spiders (Pycnogonida: Latreille, 1810) [37], and arachnids (Arachnida: Lamarck, 1801). In addition, rooted phylogenies show the most basal CFV lineages are either tick-associated (Mpulungu virus and the tick-borne CFVs) or have no known vector [36] (**Fig. 3a**). While much uncertainty remains, it is interesting to note that, if we parsimoniously assume that the basal lineages without known vectors are tick-borne (or were originally before subsequently losing their association with arachnids), phylogenies suggest (i) emergence of tick-borne viruses from arachnid-specific viruses, followed by (ii) emergence of insect-borne viruses from tick-borne viruses (**Fig. 3b**). Intriguingly, we identified synapomorphic amino acid variation in the NS5 protein, wherein ancestral residues are conserved between ancestral, arachnid-associated flaviviruses but variable in the more derived, insect-vectored flaviviruses (**Fig. 3a**, **Fig. S10**). While these patterns can of course be interpreted in alternative ways, they are consistent with positive selection accompanying the adaptation of ostensibly tick-borne viruses to newly acquired insect vectors.

## DISCUSSION

The flavivirids are a genetically and ecologically diverse group of viruses that include an unusually large number of taxonomically recognised species, many of which are associated with disease [18]. As a diverse and highly studied group, flavivirids offer unique possibilities for researchers interested in applying comparative approaches to viruses [15, 17, 18]. Species richness in this group to some extent reflects their historical importance in the development of virus research - YFV being the first human virus identified [38] - and sampling of flavivirid diversity consequently shows historical bias toward potential vector/reservoir species [39, 40]. However, with dramatic advances in DNA sequencing technology it is now possible to investigate flavivirid distribution and diversity much more broadly, building on previous comparative investigations [15–18].

In this report we address two important challenges to effective use of flavivirid sequence data in comparative genomic studies. First, we implemented our analyses using a computational framework that supports re-use of underlying data sets and facilitates reproduction of comparative genomic analyses. Second, we calibrated the long-term evolutionary history of flavivirids through use of the ‘genomic fossil record’, thereby providing broad evolutionary context for interpreting their distribution and diversity.

We identify flavivirid-derived EVEs that are >100 My old, demonstrating that the evolution of family *Flaviviridae* spans geological eras. Furthermore, we show that the robust calibrations obtained from EVEs can be combined with more tentative calibrations based on co-phyletic analysis to produce a cohesive overview of flavivirid evolution in which the major lineages emerged during the early evolution of multicellular animals and subsequently evolved together with major animal phyla (**Fig. 5c**). The timeline suggested by our analysis raises interesting questions about the ways in which animal evolution may have impacted flavivirids, since it encompasses the development of entire organ systems (e.g., the liver and vascular system) and spans the evolution of fundamental changes in animal physiology, such as the emergence of endothermy (“warm blood”) in vertebrates. The identification of flavivirid-derived EVEs in animals lacking a circulatory system (e.g., cnidarians and priapulids) suggests that cell-to-cell transmission via exosomes - as has been reported for tick-borne flaviviruses [41] - might represent the ancestral mode among flavivirids.

**Figure 5.**
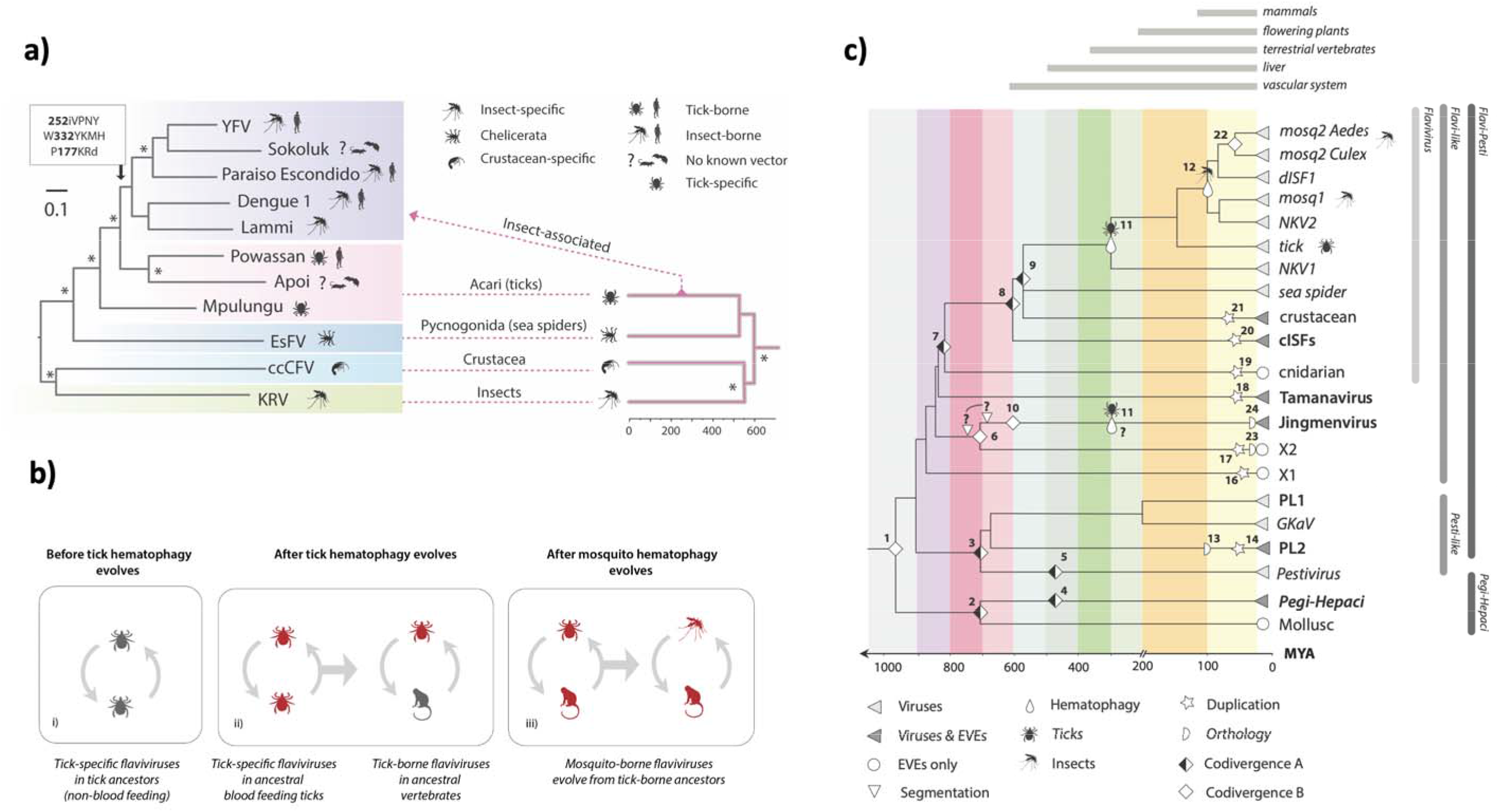
The timeline of flavivirus evolution (a) Evolution of arthropod-borne flaviviruses. The phylogeny shown on the left was constructed from an alignment spanning 804 residues of the precursor polyprotein (substitution model=LG likelihood). The phylogeny on the right is a time-calibrated phylogeny of arthropod hosts/vectors of classical flaviviruses, obtained via TimeTree [72]. The figure shows the hypothesised relationships between the host and virus trees, with virus phylogeny broadly following host phylogeny at higher taxonomic ranks but diverging dramatically from this pattern when tick-borne viruses of vertebrates emerge. **(b) Proposed model for the origin and evolution of the vector-borne flaviviruses.** Three stages are shown: (i) an ancestral group of non-vectored tick viruses is present; (ii) tick hematophagy provides prolonged exposure to vertebrate blood, thereby affording tick-specific flaviviruses the opportunity, through chance events and mutation, to acquire a capacity to replicate in vertebrate cells and ultimately become tick-borne flaviviruses of vertebrates; (iii) the existence of tick-borne flaviviruses occasions viraemic vertebrate hosts, exposing mosquitoes and other hematophagous insects to flaviviruses via blood-feeding, and ultimately allowing them to acquire a vector role. **(c) A model for long-term evolution of flavivirids.** A time-scaled tree summarising the phylogenetic relationships of flavivirids and calibrated using information obtained from analysis of endogenous viral elements (EVEs), co-phyletic analysis and the fossil record of hematophagous arthropods. Symbols on the phylogenies indicate types of calibration as shown in the key. Numbers adjacent to symbols link to rows in **Table 3** ‘Codivergence type A’ = codivergence supported by co-phylogeny. Codivergence type B = potential codivergence-based calibrations without supporting evidence. Vertical lines to right of the tree show taxonomic groups.

The vertebrate circulatory system evolved >400 Mya, and is thought to have been established in its basic form in endothermic vertebrates (i.e., birds and mammals) by ∼200 Mya [42]. Its role in transporting nutrients makes it a highly attractive target for parasitism, and hematophagous, arthropod parasites of vertebrates are thought to have evolved on >20 independent occasions [43]. Whenever this occurred, it would have created new, intimate contact networks between arthropod and vertebrate species so that viruses circulating within each group could potentially encounter opportunities to expand into the other [33]. Interestingly, we identified closely related, X2-derived EVEs in both copepods and fish, raising the possibility that copepod parasitism has enabled X2-like arthropod viruses to expand into vertebrate hosts (**Fig. 3f**).

Both phylogenetic relationships within genus *Flavivirus*, and the extended evolutionary timeline implied by our study, are consistent with a parsimonious evolutionary scenario wherein the vector-borne classical flaviviruses originate in hematophagous arachnids and later acquire the capacity to be transmitted by hematophagous insects (**Fig. 5a-b**). This ‘ticks first’ model is quite appealing because the circumstances of tick feeding provide an obvious opportunity for tick flaviviruses capable of limited replication in vertebrate cells to emerge. Feeding entails a relatively long exposure, and ticks are thought to transmit infection to one another via host blood when multiple individuals feed in proximity [44]. Once tick flaviviruses had acquired the capacity to generate sustained viremia in vertebrate hosts, opportunities for hematophagous insects to acquire vector roles would presumably arise as they become exposed to virus via blood-feeding. Interestingly, this model of flavivirus evolution is consistent with evidence that the 3’ untranslated region (UTR) of arthropod-borne flaviviruses – which can modulate the viral life cycle in complex and nuanced ways - evolved via duplication [45, 46]. Duplicated UTR sequences are best preserved in tick-borne flaviviruses, consistent with an ancestral origin, while only remnants are present in the more derived, mosquito-borne flaviviruses [47].

Due to the overall scarcity of flavivirid-derived EVEs, our investigation provides only limited insight into the evolutionary history of certain flavivirid groups, such as the *Hepaci-*, *Pegi-* and *Pestivirus* genera. However, our identification of ‘pegi-hepaci’-like EVEs in invertebrates, and the recent description of a pestivirus-like EVE in an insectivore [48], indicates that additional, diverse flavivirid EVEs will be identified as whole genome sequencing of animal species advances, allowing for more complete perspective on the long-term evolution of flavivirids. New information obtained through broader sampling of contemporary flavivirid diversity will also allow for more rigorous testing of the evolutionary hypotheses presented here.

Phylogenetic reconstructions utilising currently available data suggest that reorganisation of the flavivirid taxonomy should be considered (**Fig. 3**). Currently, family *Flaviviridae* is placed in order Amarillovirales, class Flasuviricetes, which contains no other orders or families. Flasuviricetes could be restructured to contain groups representing the ‘pegi-hepaci’ and ‘flavi-pesti’ lineages and the subgroups contained therein. This would enable taxonomic classifications to reflect the deep evolutionary splits between flavivirid subgroups, such as those separating ‘jingmenviruses’, ‘tamanaviruses’ and flaviviruses (genus *Flavivirus*) in the ‘flavi-pesti’ lineage. The enigmatic ‘Tamana bat virus’ (TABV), which was isolated in 1973 from insectivorous bats and has no known arthropod vector, has puzzled flavivirologists for decades [36, 49, 50]. It is sometimes considered a basal member of genus *Flavivirus* but is only distantly related to other flaviviruses and clearly distinct in signature genomic traits such as nucleotide composition [51]. Notably, however, we identified an EVE in the genome of the peach blossom jellyfish (*Craspedacusta sowerbii*: Lankester, 1880) that is more ‘flavivirus-like’ than TABV (**Fig. 3d**). The identification of flavivirus-like EVEs in cnidaria (Hatschek: 1888), a basal animal lineage, combined with new information describing the extensive diversity and broad host range of TABV-related viruses [8, 10, 52, 53] (**Fig. 3e**), suggests ‘tamanaviruses’ have distinct evolutionary origins to genus *Flavivirus* and should be given separate taxonomic status among ‘flavi-like’ viruses. The same applies to ‘jingmenviruses’, which presumably became evolutionarily separated from other ‘flavi-like’ viruses when they evolved multipartite genomes [11].

Given the extent of uncharacterised flavivirid diversity and the accelerating rate of virus discovery, further expansion and revision of flavivirid taxonomy will no doubt be required as exploration of the virome proceeds - numerous novel and divergent flavivirids have been described since we originally submitted this report [34, 54] - the taxonomic system proposed above would provide a relatively open structure capable of accommodating novel flavivirid diversity.

Our analysis indicates that incorporation of flavivirid-derived DNA into animal germlines is uncommon. However, in those taxa where flavivirid-derived EVEs occur, they are frequently multicopy, with some insect groups exhibiting relatively high copy number (**Table 1**). Notably, these include *Aedes* mosquitoes in which EVEs have been shown to produce PIWI-interacting RNAs (piRNAs) that limit infection with related viruses [55]. While recent data indicate a role for EVEs in antiviral defence [25, 31, 56], it remains unclear whether germline ‘capture’ of virus sequences represents a dynamic system of heritable antiviral immunity analogous to that of CRISPR-Cas. It should be noted that the distribution and diversity of cISF-derived EVEs in *Aedes* mosquito genomes is consistent with germline integration of genome-length cDNA, followed by intra-genomic amplification/fragmentation of the integrated sequences (e.g., mediated by transposable elements). Numerous, similar examples of amplification of virus-derived DNA within the animal germline have been described, involving a diverse range of virus families [57–60]. Thus, the presence of numerous distinct, flavivirid-derived EVE loci in the *Aedes* germline could reflect a relatively small number of germline colonisation events, rather than a dynamic process of EVE acquisition in association with immunity.

Controlling the spread of flavivirid-associated diseases is a public health priority and genomic data have a critical role to play in these endeavours [4, 61]. Here, we used the GLUE software framework to capture flavivirid genome sequence data and evolution-related domain knowledge in a way that supports their future use. By following principles of ‘data-oriented programming’, wherein an explicit separation is maintained between data and the code that operates on it [62], the GLUE framework can facilitate the implementation of stable data resources that make no assumptions with respect to their future usage, so they can be deployed in distinct analysis contexts [27]. Besides acting as a stable repository of domain knowledge, Flavivirid-GLUE provides a broad foundation for the rapid development of tools/services (e.g., epidemiological tracking, variant analysis) focussed on individual flavivirid taxa (e.g., see [63, 64]). In addition, Flavivirid-GLUE can underpin analytical procedures that require a broader taxonomic scope, such as sequence-based classification of newly identified flavivirids, or supporting empirical, laboratory-based investigations of important flavivirid traits (e.g., the capacity to replicate in both arthropod and vertebrate hosts). If - as our investigation suggests – flavivirid traits have been acquired gradually through long-term evolutionary interactions with animal hosts, comparative studies will likely yield many useful insights into their biology.

## MATERIALS & METHODS

### Construction of sequence-based resources for comparative genomic analysis

We used the GLUE software framework [27] to create Flavivirid-GLUE [28], an openly accessible online resource for comparative analysis of flavivirid genomes. The advantages of GLUE include: (i) the organization of virus sequence data in relation to their hypothesized evolutionary relationships (an approach that is key to the practical interpretation of sequences); (ii) integration with wider computer systems, using standard technologies such as MySQL and JSON; (iii) mechanisms to customize functionality on a usage-specific basis, for example schema extension and scripting mechanisms; (iv) features and functionality to streamline rapid development of bespoke analysis projects.

We extended GLUE’s core database schema to capture information specific to flavivirid reference sequences (e.g., virus isolate names, isolation species, date and location of sampling) and EVE loci (e.g., species in which they occur, genomic coordinates within contigs/chromosomes). A library of flavivirus representative genome sequences (**Table S1**) was obtained from GenBank via reference to the International Committee for Taxonomy of Viruses website (see [28]). GenBank sequence entries in XML format were imported into the Flavivirid-GLUE project using an appropriately configured version of GLUE’s GenBank importer module. We extracted isolate-specific information (e.g., date and location of isolation, isolate name, host species) from XML files as well standard GenBank fields (e.g., submission date). Additional/missing data was loaded from tabular files using GLUE’s TextFilePopulator module. Via reference to previous studies [6, 9, 10, 18, 53] we assigned all flavivirus sequences included in Flavivirid-GLUE to a taxonomic group and defined a standard set of genome features for flavivirids. The coordinates of genome features (where known) within all master references were recorded within the Flavivirid-GLUE database. These reference sequences and annotations were used in combination with a codon-aware, BLAST-based sequence aligner implemented in GLUE [27, 65] to generate constrained MSAs (i.e., MSAs in which the coordinate space is constrained to a selected ‘master’ reference) for each taxonomic rank within the *Flaviviridae* (**Table 1**). To address genome and gene-coverage-related issues we used GLUE to generate genome feature coverage data for member sequence. Constrained MSAs were used to infer the coordinates of genome features not explicitly defined in GenBank XML via GLUE’s ‘inherit features’ command [27].

### Genome screening in silico

Systematic *in silico* genome screening was performed using the database-integrated genome screening (DIGS) tool [66] – a PERL-based screening framework within which the basic local alignment search tool (BLAST) program suite [65] is used to perform similarity searches while the MySQL relational database management system (Community Server 8.0.26) is used to record their output. WGS data were obtained from the National Center for Biotechnology Information (NCBI) genome database [67] - we obtained all animal genomes available as of March 2020 (see [28]). Flavivirid reference genomes and coding feature annotations collated in Flavivirid-GLUE were used to derive polypeptide probes for tBLASTn-based screening in the DIGS framework. For virus genomes lacking detailed annotations, we created polypeptide probes based on fragments of the major polyprotein. Via screening of WGS assemblies using the DIGS tool we generated a non-redundant database of flavivirid-derived EVE loci [28]. We used DIGS to investigate these loci and categorise them into: (i) putatively novel EFV elements; (ii) orthologs of previously characterised EFVs (e.g., copies containing large indels); (iii) non-viral sequences that cross-matched to flavivirus probes (e.g., retrotransposons). In applying IDs to EVE sequences (see **Fig. 2**, **Table 1**), we conservatively assumed that the presence of multiple EVE loci generally reflects intragenomic amplification rather than multiple independent germline incorporation events.

### Phylogenetic and genomic analysis

Flavivirid-GLUE was used to implement an automated process for reconstructing midpoint-rooted, bootstrapped phylogenies from MSA partitions representing each rank within the constrained MSA tree. Gene coverage data was used to condition the way in which taxa were selected into MSA partitions [68]. Phylogenies were reconstructed using the maximum likelihood approach implemented in RAxML (version 8.2.12) [68]. Protein substitution models were selected via hierarchical maximum likelihood ratio test using the PROTAUTOGAMMA option in RAxML. JalView [69] (version 2.11.1.4) and Se-Al (version 2.0a11) were used to inspect MSAs.

## Data Availability

Data available via GitHub: https://giffordlabcvr.github.io/Flavivirus-GLUE/

## Acknowledgements

RJG was funded by the Medical Research Council of the United Kingdom (MC_UU_12014/12). WMS acknowledges support from the Global Virus Network Fellowship. We thank Anna Gatseva, Joseph Hughes, Alain Kohl, Spyros Lytras, Emilie Pondeville, Charles Rice, David Robertson, Greg Towers, Sam J. Wilson, and anonymous reviewers for critical reading of the manuscript.

