## Supplementary figures for "Comparative analysis of genome-encoded viral sequences reveals the evolutionary history of flavivirids (family *Flaviviridae*)"

Fig. S1

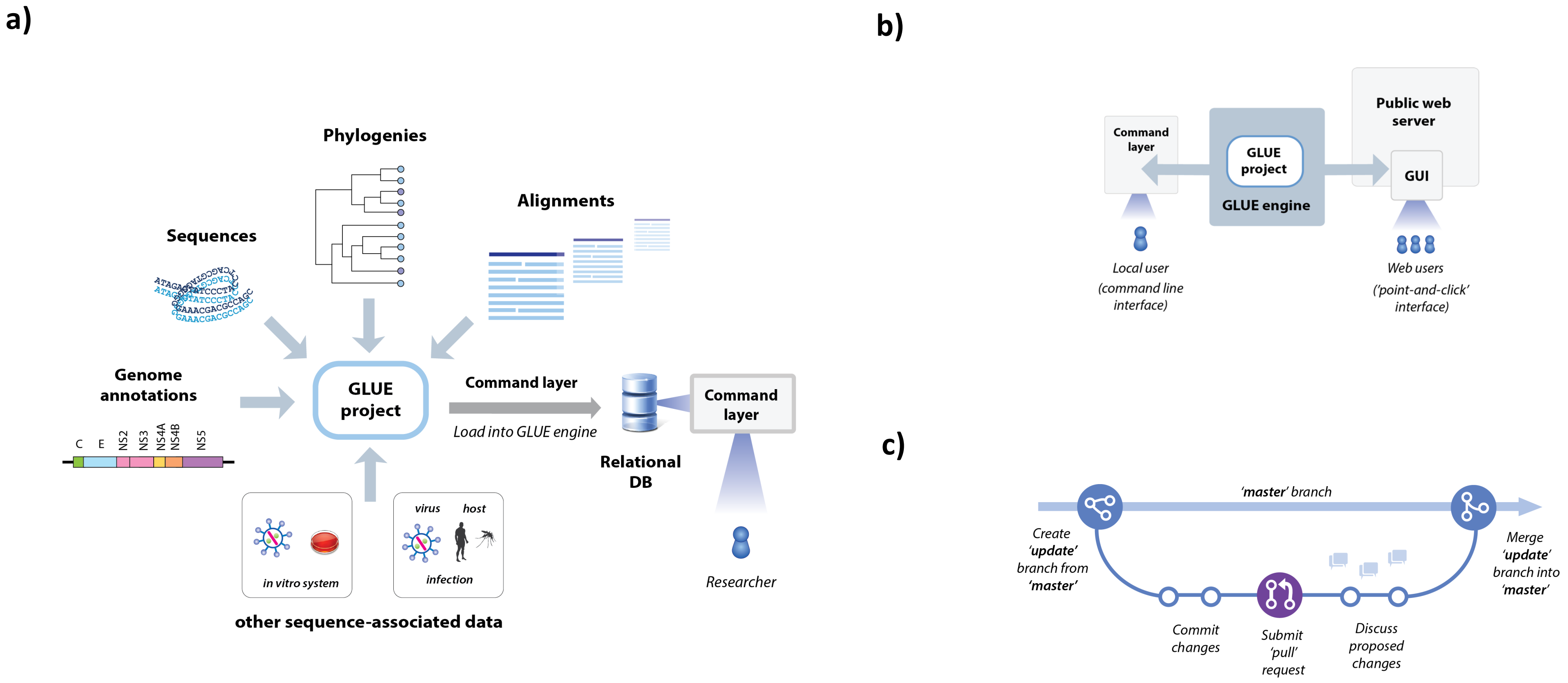
a)
Phylogenies
Alignments
Sequences
Genome annotations
C
E
NS2
NS3
NS4A
NS4B
NS5
GLUE project
Command layer
Load into GLUE engine
Relational DB
Command layer
in vitro system
virus
host
infection
Researcher
other sequence-associated data
b)
Public web server
Command layer
GLUE project
GLUE engine
GUI
Local user (command line interface)
Web users ('point-and-click' interface)
c)
'master' branch
Create 'update' branch from 'master'
Merge 'update' branch into 'master'
Commit changes
Submit 'pull' request
Discuss proposed changes

**Fig. S2**

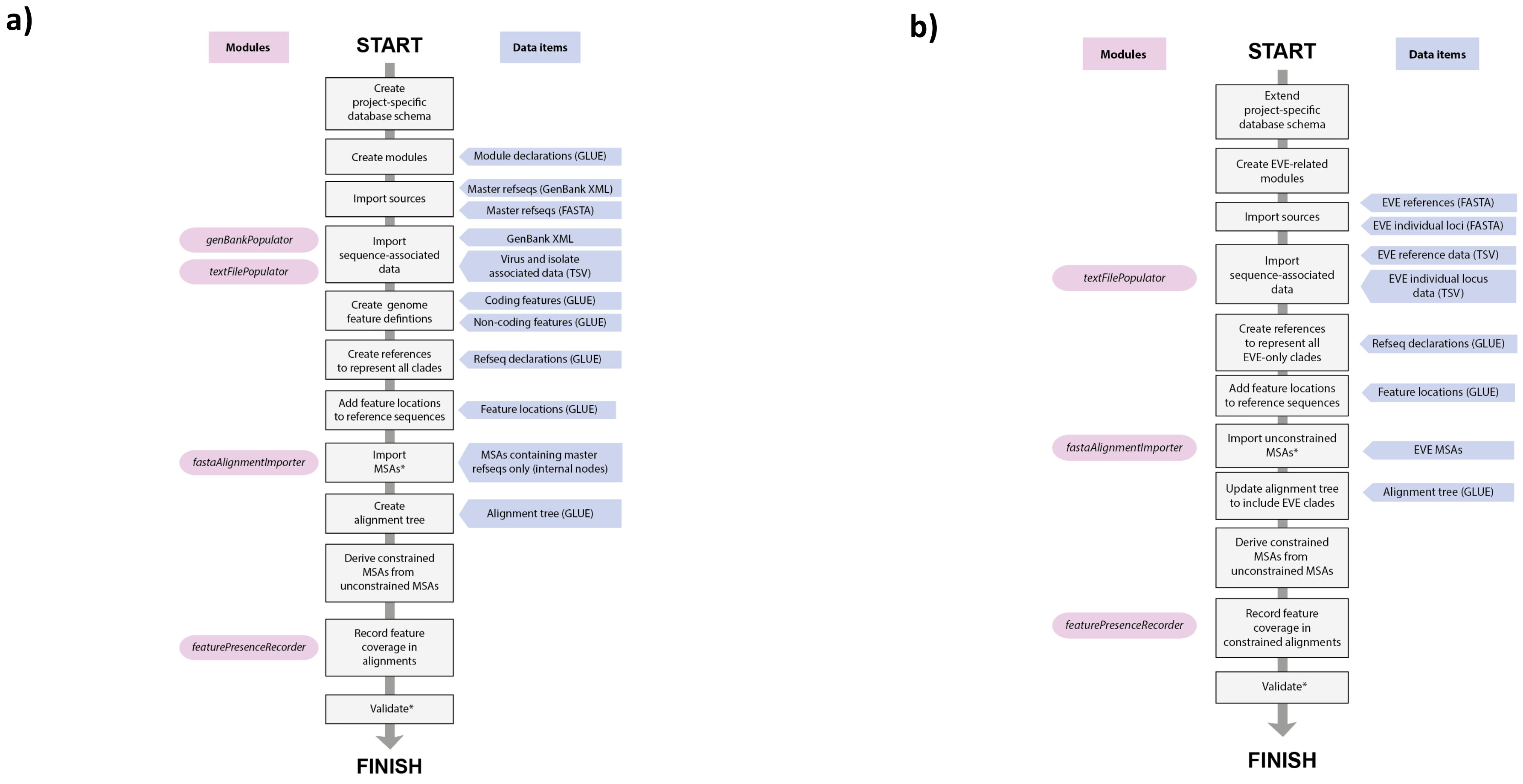
a)
Modules
START
Data items
Create project-specific database schema
Create modules
Module declarations (GLUE)
Import sources
Master refseqs (GenBank XML)
Master refseqs (FASTA)
genBankPopulator
textFilePopulator
Import sequence-associated data
GenBank XML
Virus and isolate associated data (TSV)
Create genome feature defintions
Coding features (GLUE)
Non-coding features (GLUE)
Create references to represent all clades
Refseq declarations (GLUE)
Add feature locations to reference sequences
Feature locations (GLUE)
fastaAlignmentImporter
Import MSAs*
MSAs containing master refseqs only (internal nodes)
Create alignment tree
Alignment tree (GLUE)
Derive constrained MSAs from unconstrained MSAs
featurePresenceRecorder
Record feature coverage in alignments
Validate*
FINISH
b)
Modules
START
Data items
Extend project-specific database schema
Create EVE-related modules
Import sources
EVE references (FASTA)
EVE individual loci (FASTA)
textFilePopulator
Import sequence-associated data
EVE reference data (TSV)
EVE individual locus data (TSV)
Create references to represent all EVE-only clades
Refseq declarations (GLUE)
Add feature locations to reference sequences
Feature locations (GLUE)
fastaAlignmentImporter
Import unconstrained MSAs*
EVE MSAs
Update alignment tree to include EVE clades
Alignment tree (GLUE)
Derive constrained MSAs from unconstrained MSAs
featurePresenceRecorder
Record feature coverage in constrained alignments
Validate*
FINISH

### Fig S3

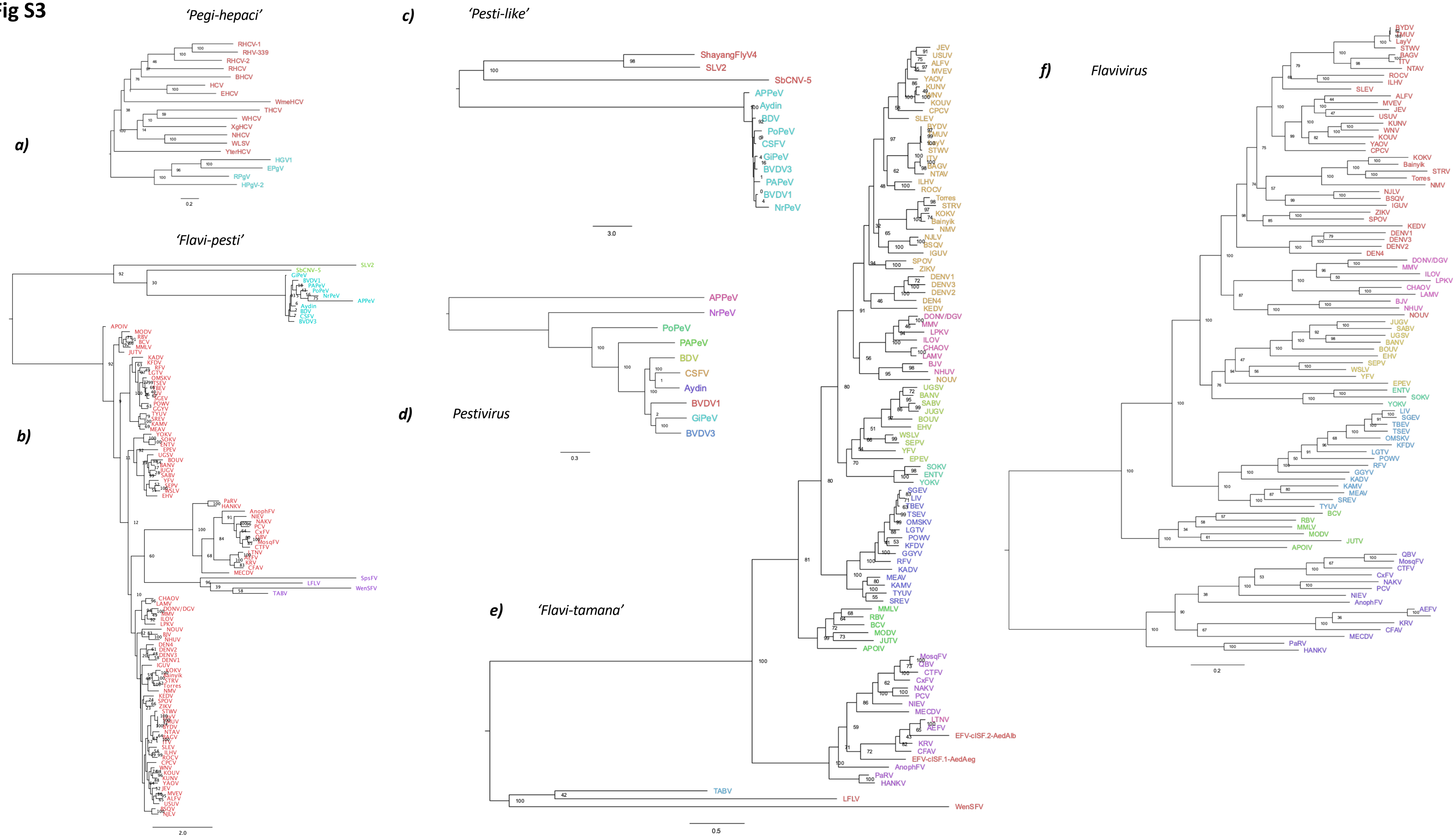
a)
'Pegi-hepaci'
b)
'Flavi-pesti'
c)
'Pesti-like'
ShayangFlyV4
SLV2
SbCNV-5
APPeV
Aydin
BDV
PoPeV
CSFV
GiPeV
BVDV3
PAPeV
BVDV1
NrPeV
d)
Pestivirus
APPeV
NrPeV
PoPeV
PAPeV
BDV
CSFV
Aydin
BVDV1
GiPeV
BVDV3
e)
'Flavi-tamana'
f)
Flavivirus

**Fig. S4**

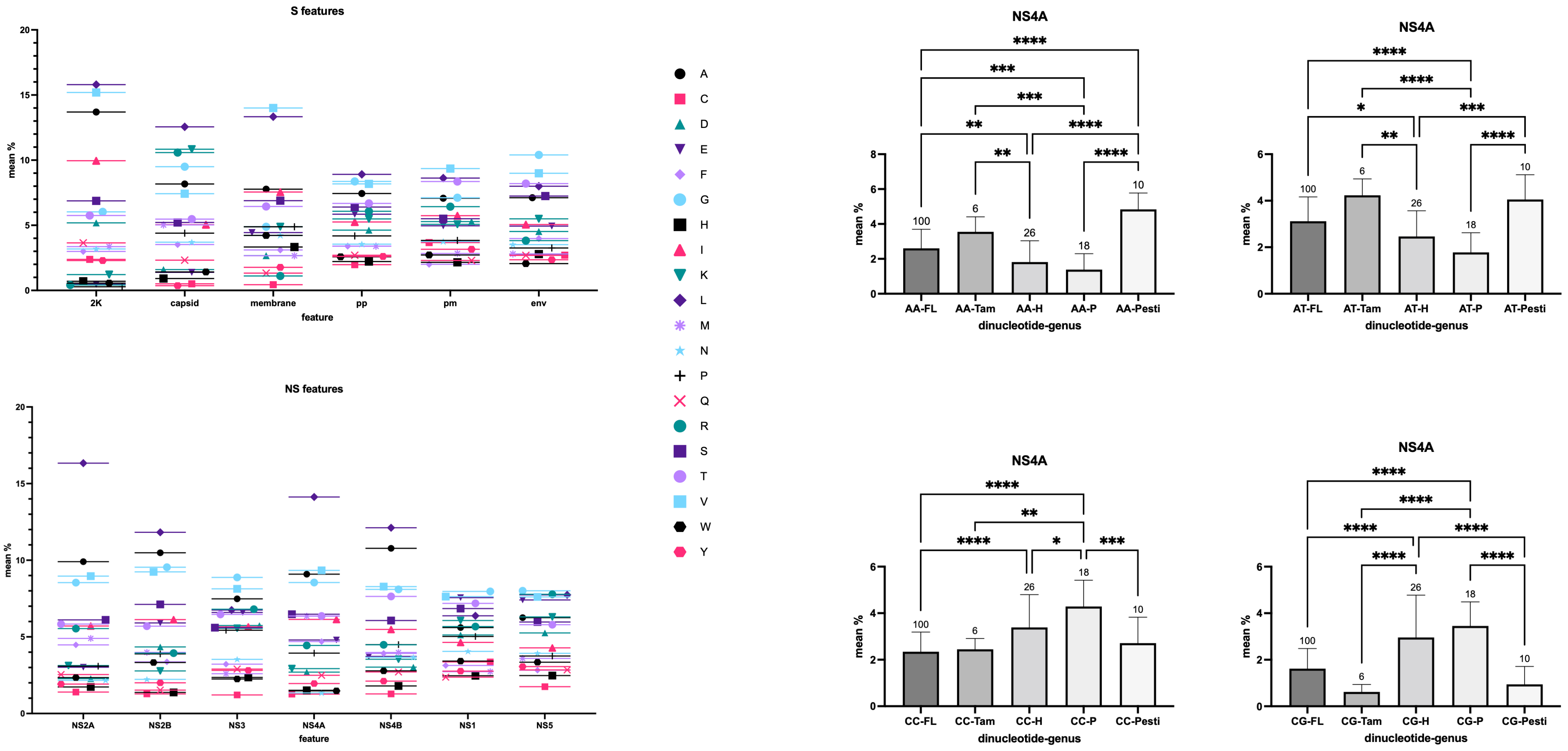
S features
mean %
2K
capsid
membrane
pp
pm
env
feature
NS features
mean %
NS2A
NS2B
NS3
NS4A
NS4B
NS1
NS5
feature
A
C
D
E
F
G
H
I
K
L
M
N
P
Q
R
S
T
V
W
Y
NS4A
mean %
AA-FL
AA-Tam
AA-H
AA-P
AA-Pesti
dinucleotide-genus
NS4A
mean %
AT-FL
AT-Tam
AT-H
AT-P
AT-Pesti
dinucleotide-genus
NS4A
mean %
CC-FL
CC-Tam
CC-H
CC-P
CC-Pesti
dinucleotide-genus
NS4A
mean %
CG-FL
CG-Tam
CG-H
CG-P
CG-Pesti
dinucleotide-genus

**Fig. S5**

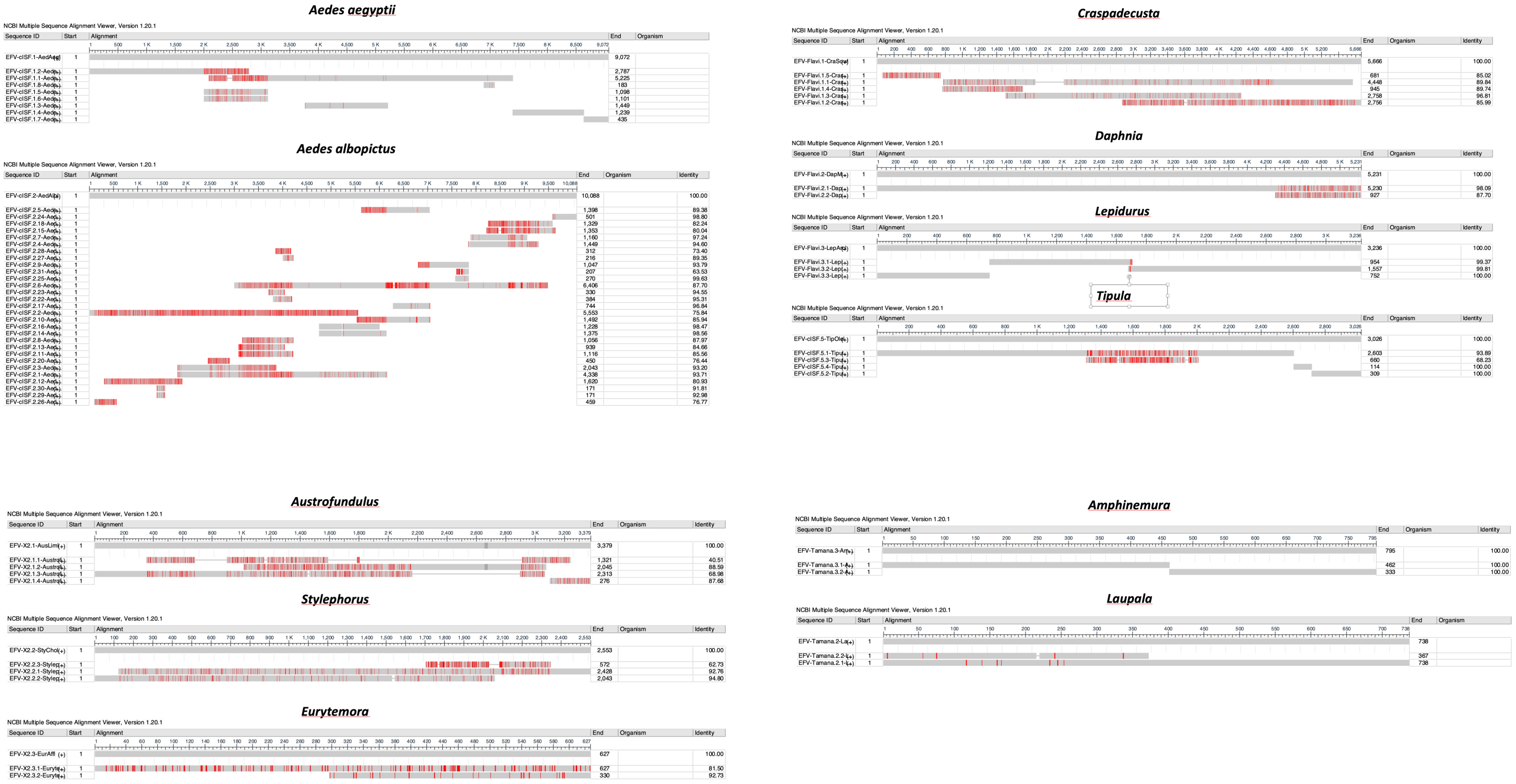
Aedes aegyptii
Aedes albopictus
Craspadecusta
Daphnia
Lepidurus
Tipula
Austrofundulus
Stylephorus
Eurytemora
Amphinemura
Laupala
NCBI Multiple Sequence Alignment Viewer, Version 1.20.1
Sequence ID
Start
Alignment
End
Organism
Identity

### Fig. S6

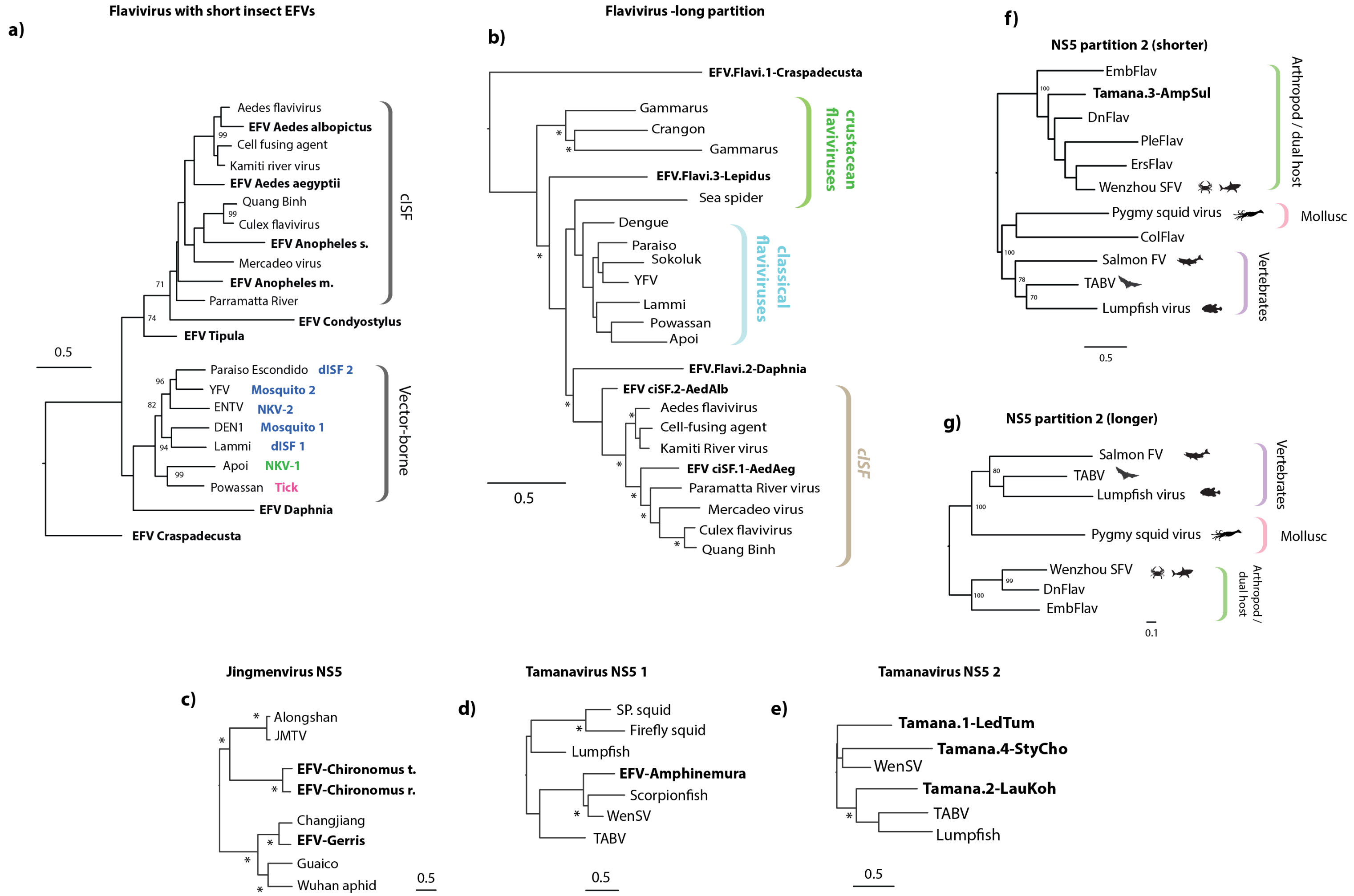

a)
Flavivirus with short insect EFVs
Aedes flavivirus
EFV Aedes albopictus
Cell fusing agent
Kamiti river virus
EFV Aedes aegyptii
Quang Binh
Culex flavivirus
EFV Anopheles s.
Mercadeo virus
EFV Anopheles m.
Parramatta River
EFV Condyostylus
EFV Tipula
cISF
Paraiso Escondido dISF 2
YFV Mosquito 2
ENTV NKV-2
DEN1 Mosquito 1
Lammi dISF 1
Apoi NKV-1
Powassan Tick
EFV Daphnia
EFV Craspadecusta
Vector-borne
0.5
b)
Flavivirus -long partition
EFV.Flavi.1-Craspadecusta
Gammarus
Crangon
Gammarus
EFV.Flavi.3-Lepidus
Sea spider
crustacean flaviviruses
Dengue
Paraiso
Sokoluk
YFV
Lammi
Powassan
Apoi
classical flaviviruses
EFV.Flavi.2-Daphnia
EFV ciSF.2-AedAlb
Aedes flavivirus
Cell-fusing agent
Kamiti River virus
EFV ciSF.1-AedAeg
Paramatta River virus
Mercadeo virus
Culex flavivirus
Quang Binh
cISF
0.5
f)
NS5 partition 2 (shorter)
EmbFlav
Tamana.3-AmpSul
DnFlav
PleFlav
ErsFlav
Wenzhou SFV
Arthropod / dual host
Pygmy squid virus
Mollusc
ColFlav
Salmon FV
TABV
Lumpfish virus
Vertebrates
0.5
g)
NS5 partition 2 (longer)
Salmon FV
TABV
Lumpfish virus
Vertebrates
Pygmy squid virus
Mollusc
Wenzhou SFV
DnFlav
EmbFlav
Arthropod / dual host
0.1
c)
Jingmenvirus NS5
Alongshan
JMTV
EFV-Chironomus t.
EFV-Chironomus r.
Changjiang
EFV-Gerris
Guaico
Wuhan aphid
0.5
d)
Tamanavirus NS5 1
SP. squid
Firefly squid
Lumpfish
EFV-Amphinemura
Scorpionfish
WenSV
TABV
0.5
e)
Tamanavirus NS5 2
Tamana.1-LedTum
Tamana.4-StyCho
WenSV
Tamana.2-LauKoh
TABV
Lumpfish
0.5

**Fig. S7**

a)

```
Ceratina-LSNX01048950 ATATAATCAGCCCCCGGGACTTTGGTAGTGTTTCTCATTTGTTGAGTGTGAACTATTGAA
Euglossa-NIJG01001146 TTATAATGA------GGCGATTTGATTGTGATAATCAAATGATAATCAAATGATAATCAA
                       ****** *      **   **** * *** *  ***  ** * *        ** * **
Ceratina-LSNX01048950 GAATTTTTCCTCAGGGTTTTGCCGTAGGCGCATCCGAGTGTATCGGACAATAATAAGTTG
Euglossa-NIJG01001146 AATGTTTTCCTGCGGATTTCCGTATTTGCGTTTCTGAGATCATCTGACATTAA-AAGCTT
                          *******  ** ***     *  ***  ** ***   *** **** *** *** *
Ceratina-LSNX01048950 AATCAAACGTTGAATAATAGCCTCGTTTGTTTCGCTTACTTAATTTAAGTTTAAAATAAG
Euglossa-NIJG01001146 AATCTACTATTCAGGCGTTACC-CGCTTGGTGGGGGT--TTAGTTTGAGTTT------GG
                      **** *   ** *    *  ** ** *** *  *     *** *** *****       *
Ceratina-LSNX01048950 AGCCATTAGCTCTTAATTATTATT---------ATTTCTATTAATTTAG-TTTAGTAAT-
Euglossa-NIJG01001146 GGCGATTGGCCTCATTTGAATATTTAAGCTTAAATATTTTTCAATTTAATTTTAGTGTTT
                       ** *** **      * * ****         ** * * * ******  ****** *
Ceratina-LSNX01048950 -ATTACTTAAATATTA-TAATCAGTAATATAAGTCATATCTTACTTAGTAATTTAGTTCG
Euglossa-NIJG01001146 AGTTACTTATTTAGCGTTACTTAGCAAATTAAGGAATGACATAACT-TTAAGTTAATTCG
                        *******  **    ** * ** **  ****  **    **  *   ** *** ****
Ceratina-LSNX01048950 ---TATAGGTATAAGGGTTCTTTATTCTTTTGTTTACTAGTTTAGTGGTAGACGC-ATAA
Euglossa-NIJG01001146 TCGTTGCGACTTATAGATTAAGGATAGGTAAATTAAATAGTTAATAGGTGTTCTCAATAA
                         *   *   **  * **    **   *   ** * ***** *  ***   * * ****
ORF                                                                     M  S  S
Ceratina-LSNX01048950 GAT--TTTTGACTAGTATAATATTCTTGATTATTTTTAAAAAGGTTTCAGCATGTCGTCC
Euglossa-NIJG01001146 GATAACTTAAGTTTAAATAATAGTATCATTAGTTTTTAAAAAGGTTTCAGCATGTCGTCC
                      ***   **    *   ******** *   *  *******************
ORF                    Q  N  I  S  R  Q  K  I  G  A  I  R  S  Q  A  A  S  A  E  V
Ceratina-LSNX01048950 CAGAATATCTCTAGGCAGAAAATTGGGGCGATTCGCTCGCAAGCTGCTTCGGCTGAAGTA
Euglossa-NIJG01001146 CAGAATATCTCTAGGCAGAAAATTGGGGCGATTCGCTCGCAAGCTGCTTCGGCTGAAGTA

ORF                    A  A  Q  M  R  A  V  A  A  K  A  E  E  V  A  K  L  A  E  K
Ceratina-LSNX01048950 GCGGCTCAGATGAGAGCAGTGGCGGCCAAAGCAGAAGAAGTTGCTAAATTGGCTGAAAAG
Euglossa-NIJG01001146 GCGGCTCAGATGAGAGCAGTGGCGGCCAAAGCAGAAGAAGTTGCTAAATTGGCTGAAAAG

ORF                    G  V  D  E  Q  R  Q  V  L  A  N  A  G  V  A  P  P  A  V  L
Ceratina-LSNX01048950 GGCGTGGACGAACAGCGGCAAGTACTTGCAAATGCTGGCGTTGCTCCGCCTGCTGTTTTG
Euglossa-NIJG01001146 GGCGTGGACGAACAGCGTGAAGTGTTGCAACCAGATGGAATTGCACCACCAGTGGTTCTT

ORF                    N  R  K  K  R  T  V  K  K  P  K  G  L  L  Q  R  V  M  R  D
Ceratina-LSNX01048950 AATCGTAAGAAGAGAACAGTTAAAAAACCGAAAGGATTGCTTCAGCGGGTCATGAGGGAT
Euglossa-NIJG01001146 AATCGAAAGAAAAGGCGTTCATGACACACGAAGGGTTTGGGTACAGATGACATTGGAATC

ORF                    ?  L  V  R  L  C  R  *  F  A  *  P  C  C  S  Q  R  A  W  E
Ceratina-LSNX01048950 --GCTAGTACGTTTATGCAGATAATTTGCATAACCTTGTTGCTCACAACGTGCATGGGAG
Euglossa-NIJG01001146 -AGCTAGTAAGTTAGTTTCGT------------------TTGTTTGTGTGTGCATGGCTT

ORF                    L  K  F  R  V  A  T  R  D  E  K  A  R  F  V  Y  *  I  E  T
Ceratina-LSNX01048950 CTCAAGTTCCGAGTGGCAACCCGGGACGAAAAAGCGAGATTTGTGTATTAGATCGAAACC
Euglossa-NIJG01001146 TGGGCTTGGCCAAACACTCCTGTACCTTTGTGTTCATTGGACGGCAGCGGTCTTAATTTG

ORF                    K  L  T  *  I  S  S  W  L  S  S  S  L  ?  I  S  D  L  S  G
Ceratina-LSNX01048950 AAATTGACCTGAATCAGCAGCTGGTTAAGCAGCTCATTGGC-ATCAGTGATCTTTCAGGC
Euglossa-NIJG01001146 CAACGA------A--CTCTGCTGGTTAAGCTTTCTAAAC---TTAACTAGCTTTACCGAC

ORF                    V  E  V  F  R  I  A  D  L  G  K  R  K  I  E  E  L  K  S  L
Ceratina-LSNX01048950 GTCGAAGTTTTTCGCATAGCCGACTTAGGGAAACGCAAGATTGAAGAGTTAAAATCTCTT
Euglossa-NIJG01001146 TTTAAGGTGGGTAAGCTAGCAGGATTGGAAGAAATGATGTCTTCCAAATTGAGAGAACTT

ORF                    G  K  D  Y  E  N  F  Y  K  A  L  S  Y  C  K  I  S  N  A  D
Ceratina-LSNX01048950 GGGAAAGATTACGAAAATTTCTATAAGGCACTTAGCTATTGTAAAATTTCAAATGCAGAC
Euglossa-NIJG01001146 CGATTAGAGTGGAACGACGTTGATAAAGCATTTAAGTATTGTCGCGTGCCGTCTGTCGAG

ORF                    V  S  K  V  G  N  L  K  Q  I  S  R  G  N  V  I  D  I  N  N
Ceratina-LSNX01048950 GTTAGTAAGGTCGGCAACTTGAAACAAATTTCTCGTGGCAATGTGATCGACATTAACAAC
Euglossa-NIJG01001146 ACTAACGTGCTGGCTGAAGTTAGACGTATTGCACGTAACAATGTCGGCAAAATCAATGGA

ORF                    V  A  S  Q  L  D  S  C  K  S  E  Y  N  R  L  L  D  V  A  K
Ceratina-LSNX01048950 GTGGCTAGTCAGTTAGACAGTTGTAAATCTGAGTACAATCGGTTGTTAGACGTGGCTAAG
Euglossa-NIJG01001146 GTATTGTCGAGTTTGGTATCCTGTCGGGAAAGCTACAAGCGGCAAGCAGAAATGCTTAAC

ORF                    T  Q  N  T  K  L  D  S
Ceratina-LSNX01048950 ACACAAAATACTAAGTTGGATTCT
Euglossa-NIJG01001146 CCTTTTAGCACCGAACGAAAGGAG
```

b)

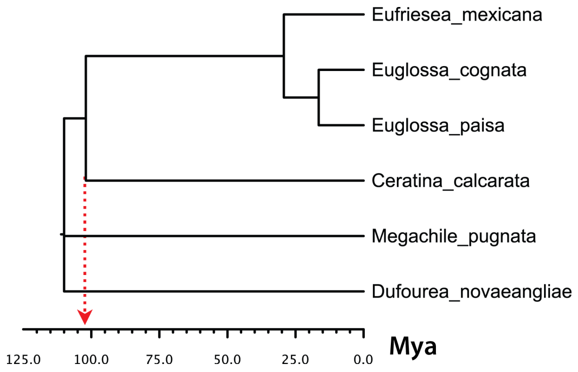

**Fig. S8**

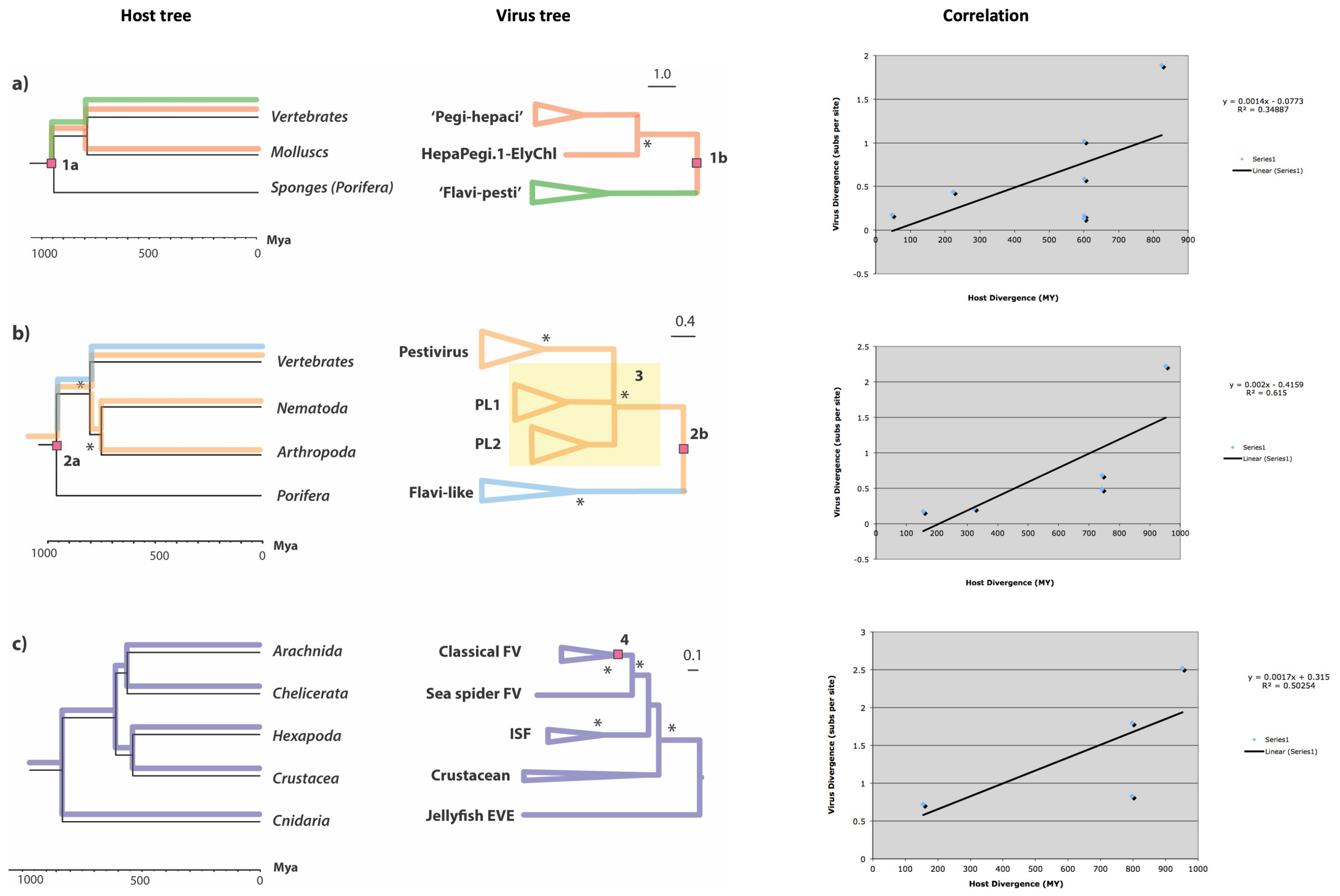

Host tree
Virus tree
Correlation
a)
Vertebrates
Molluscs
Sponges (Porifera)
1a
1000
500
0
Mya
'Pegi-hepaci'
HepaPegi.1-ElyChl
'Flavi-pesti'
1.0
1b
y = 0.0014x - 0.0773
R² = 0.34887
Series1
Linear (Series1)
Virus Divergence (subs per site)
Host Divergence (MY)
b)
Vertebrates
Nematoda
Arthropoda
Porifera
2a
Pestivirus
PL1
PL2
Flavi-like
0.4
3
2b
y = 0.002x - 0.4159
R² = 0.615
c)
Arachnida
Chelicerata
Hexapoda
Crustacea
Cnidaria
Classical FV
Sea spider FV
ISF
Crustacean
Jellyfish EVE
4
0.1
y = 0.0017x + 0.315
R² = 0.50254

**Fig. S9**

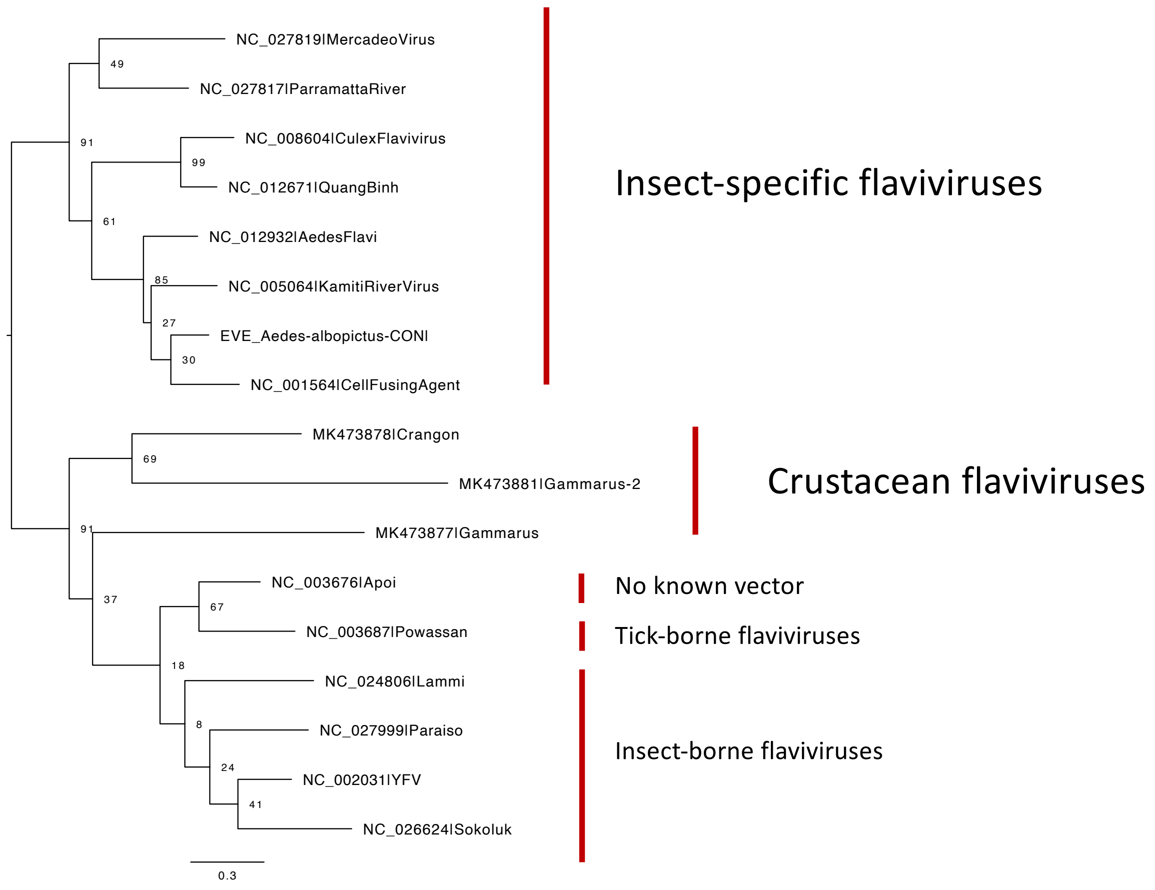
NC_027819|MercadeoVirus
49
NC_027817|ParramattaRiver
91
NC_008604|CulexFlavivirus
99
NC_012671|QuangBinh
61
NC_012932|AedesFlavi
85
NC_005064|KamitiRiverVirus
27
EVE_Aedes-albopictus-CONI
30
NC_001564|CellFusingAgent
MK473878|Crangon
69
MK473881|Gammarus-2
91
MK473877|Gammarus
NC_003676|Apoi
37
67
NC_003687|Powassan
18
NC_024806|Lammi
8
NC_027999|Paraiso
24
NC_002031|YFV
41
NC_026624|Sokoluk
0.3
Insect-specific flaviviruses
Crustacean flaviviruses
No known vector
Tick-borne flaviviruses
Insect-borne flaviviruses

### Fig. S10

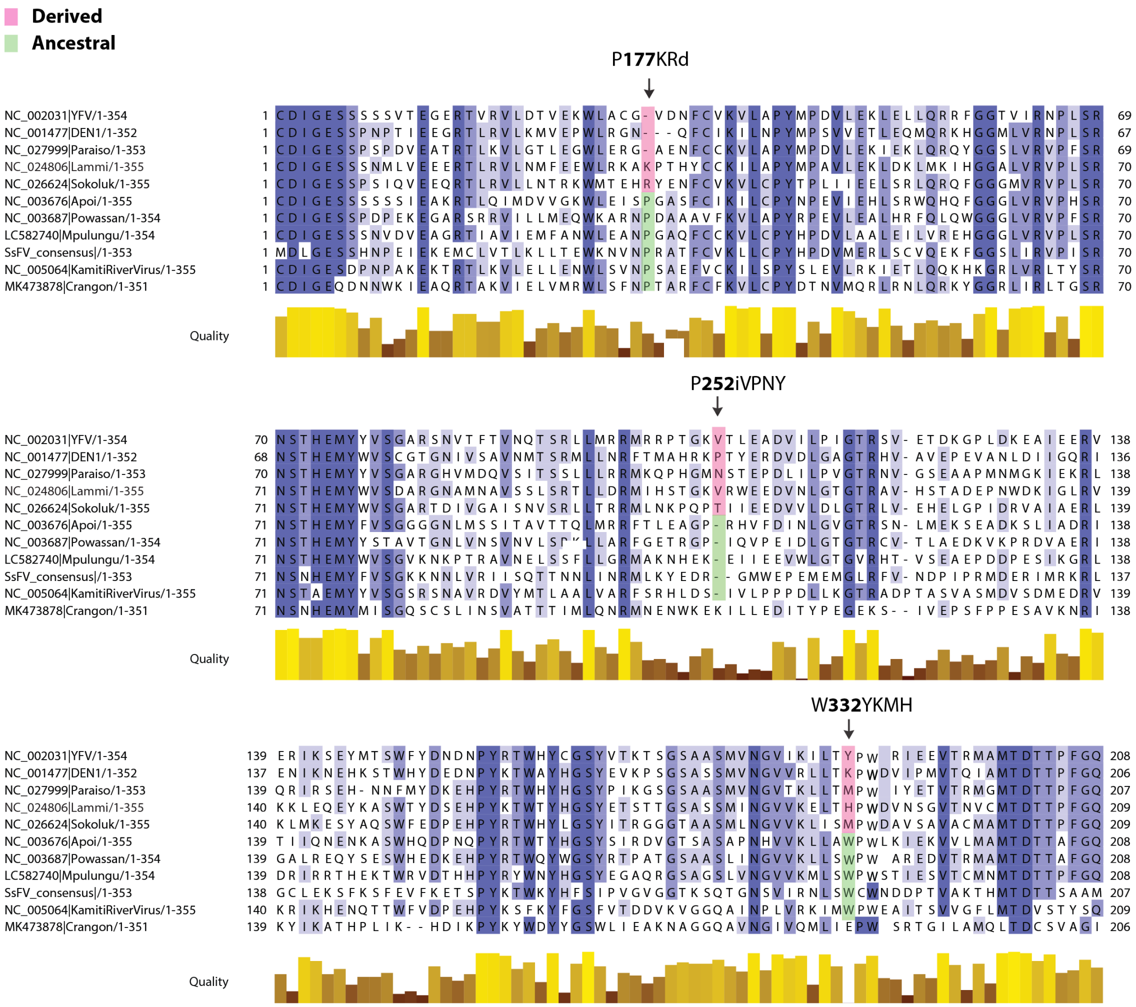
