## Supplementary figure legends for "Comparative analysis of genome-encoded viral sequences reveals the evolutionary history of flavivirids (family *Flaviviridae*)"

**Figure S1. Flavivirid-GLUE - a data-oriented approach to comparative genomic analysis of flavivirids.** The GLUE software framework can be used to develop sequence data-oriented ‘projects’ that contain molecular sequence data along with diverse additional forms of sequence-associated data (e.g., genome annotations, multiple sequence alignments and phylogenies), as shown schematically in panel (**a**). Loading projects into the GLUE ‘engine’ creates a relational database that captures the semantic relationships between data items. The database is constructed using GLUE’s native command layer, which can also be used to develop analysis protocols that utilise data items by interacting with the database and with commonly used bioinformatics programs (e.g., BLAST, RAxML, MAAFT). **(b)** By hosting on a public web server and adding a graphical user interface (GUI), projects can be developed into accessible, web-based resources that use the underlying data to provide specific services (e.g., drug resistance interpretation). **(c)** Hosting of GLUE projects in an online version control system (e.g., GitHub) allows for their controlled collaborative development.

**Figure S2. The Flavivirid-GLUE resource build process.**

Flowchart showing the process through which (i) the core project database is constructed in the Flavivirid-GLUE resource (left); (ii) the Flavivirid-GLUE project database is extended through the addition of an EVE-focussed project layer.

**Figure S3. Phylogeny reconstruction via Flavivirid-GLUE**

Bootstrapped, maximum likelihood phylogenies reconstructed for a range of taxonomic ranks within family *Flaviviridae* via GLUE, using MSAs spanning the relatively conserved NS5 gene. Phylogenies shown are as follows: (**a**) ‘Pegi-hepaci’ lineage, taxa labels coloured by virus genus; (**b**) ‘Flavi-pesti’ lineage coloured by virus genus; (**c**) ‘Pesti-like’ lineage, taxa labels coloured by virus subgroup; (**d**) *Pestivirus* genus, taxa labels coloured by virus host species; (**d**) ‘Flavi-tamana’ lineage, taxa labels coloured by virus subgroup; (**d**) *Flavivirus* genus, taxa labels coloured by virus subgroup. Phylogenies were constructed using amino acid sequences (obtained via translation of an in-frame, codon-level alignment). Scale bars indicate evolutionary distance in substitutions per site. Numbers on nodes show bootstrap support (1000 replicates). All trees are midpoint rooted for display purposes.

**Figure S4. Amino acid and dinucleotide composition analysis via Flavivirid-GLUE**

**(a)** Flavivirid-GLUE was used to obtain the % of amino acids in each feature, normalised to feature length. Structural features (S features; top) and Non-structural features (NS features; bottom) in the *Flavivirus* genus. Means and standard deviation calculated and plotted with GraphPad Prism (V9.3.0). **(b)** Dinucleotide counts in NS4A feature in five genera/groups (*Flavivirus*, *Pestivirus*, *Hepacivirus*, *Pegivirus*, and ‘tamanavirus’). The number of dinucleotides in each feature for each genus was counted and normalised to the length of the features, using GLUE, and represented as %. All dinucleotide pairs were counted, however here only dinucleotides AA, AT, CC, and CG are shown, for simplicity. The number of sequences used per genera is shown above each bar. The differences between the means of each dinucleotide pair were compared using a one-way ANOVA, only statistically significant differences are shown (*p<0.05, **p<0.01, ***p<0.001, ****p<0.0001). FL=Flavi; Tam=Tamana; H=Hepaci; P=Pegi.

**Figure S5. Structure of multicopy EVEs.**

Schematic diagrams showing alignments of multicopy EFV from the same species or species group. Figures were generated using NCBI’s alignment viewer (<https://www.ncbi.nlm.nih.gov/projects/msaviewer/>).

**Figure S6. Phylogenies showing evolutionary relationships of EFVs.**

Bootstrapped maximum likelihood phylogenies (1000 replicates), reconstructed for viruses and flavivirid-derived EVEs at different taxonomic levels, and using alternative partitions to accommodate short EVE sequences. Phylogenies were constructed using amino acid sequences (obtained via translation of a codon-level alignment). Scale bars indicate evolutionary distance in substitutions per site. Brackets to the right indicate genera and sub-lineages. Asterisks indicate bootstrap support ≥70% (1000 replicates). All trees are midpoint rooted for display purposes.

**Figure S7. Homologous flanks of PL2-Apoidae insertions.**

**(a)** An interleaved, pairwise alignment between orthologous EPVs identified in insect species the spurred ceratina (*Ceratina calcarata*: Robertson, 1900) and the green orchid bee (*Euglossa* *dilemma*: Bembé & Eltz, 2011). Taxa names and GenBank accession numbers of whole genome shotgun contigs are shown to the left of alignment rows. The translation of the open reading frame is shown on brown background above the nucleotide sequence. **(b)** A phylogeny of the superfamily Apoidae showing the estimated divergence times of included in our analysis, and the inferred minimum time since the virus that gave rise to the PL2-Apoidae lineage of endogenous viral elements was incorporated into the Apoidae germline (dashed arrow). Calibrated phylogeny was obtained via TimeTree {Kumar, 2017 #214}. MYA=millions of years ago

**Figure S8. Putative codivergence of flavivirid groups and host phyla.**

‘Tanglegrams’ illustrating matching topologies in animal (left) and flavivirid (right) phylogenies. Putative tracking of host lineages by viral lineages is indicated on the host phylogeny. Clades within which the branch lengths of virus phylogenies are correlated with divergence times in host animal lineages, as shown in the plots (right): the invertebrate and vertebrate splits in the (**a**) ‘pegi-hepaci’ lineage (R^2^=0.5); (**b**) ‘pesti-like’ lineage (R^2^=0.7); (**c**) the cnidarian-arthropod, chelicerate-hexapoda, and crustacean-insect splits in the *Flavivirus* genus (R^2^=0.3). Note that, due to limited data, all comparisons rely on strong assumptions regarding virus phylogenies, indicated in the figure by numbers, as follows: (1) and (2) midpoint rooting was used, and the deepest divergence in the virus tree was approximately calibrated using the divergence date of porifera (the most basal animal lineage) in line our hypothesis that major flavivirid lineages originated in early metazoans; (3) the splits between nematode and arthropod viruses in “pesti-like 1” (PL1) and “pesti-like 2” (PL2) clades – indicated by the yellow square - were poorly resolved in our phylogenies; (4) midpoint rooting was used, and it is assumed that the arthropod-borne classical flaviviruses originated in arachnids, as proposed in **Fig 5a**. Abbreviations: Mya=Million years ago.

**Figure S9. NS3-based reconstruction of evolutionary relationships in genus *Flavivirus***

A bootstrapped maximum likelihood phylogeny (1000 replicates), reconstructed for genus flavivirus from an alignment of conserved residues within the NS3 gene of flaviviruses (genus *Flavivirus*). Scale bars indicate evolutionary distance in substitutions per site. Brackets to the right indicate genera and sub-lineages. Asterisks indicate bootstrap support ≥70% (1000 replicates). All trees are midpoint rooted for display purposes.

**Figure S10.** An interleaved multiple sequence alignment showing homology between sea spider Flavivirid and the vector-borne, ‘classical’ Flavivirides, including representatives of the following major subclades: (i) mosquito-borne Flavivirides - Dengue virus (DEN1), Yellow fever virus (YFV); (ii) Tick borne Flavivirides – Powassan virus; (iii) no-known vector (NKV) – Apoi virus. Synapomorphies indicating ancestry of sea spider Flavivirid to the tick-borne/NKV group are highlighted by arrows. GenBank accession numbers are indicated for each taxon in the alignment. Histograms beneath each row indicate sequence conservation for each alignment column. Alignment coordinates are indicated at either side of each alignment row. Ancestral versus derived variation is highlighted as shown in the key.
